# The direct and indirect pathways of the basal ganglia antagonistically influence cortical activity and perceptual decisions

**DOI:** 10.1101/2022.08.26.505381

**Authors:** Enny H. van Beest, Mohammed A.O. Abdelwahab, J. Leonie Cazemier, Chrysiida Baltira, M. Cassandra Maes, Brandon D. Peri, Matthew W. Self, Ingo Willuhn, Pieter R. Roelfsema

## Abstract

The striatum, input nucleus of the basal ganglia, receives topographically organized input from the cortex and gives rise to the direct and indirect pathways with antagonistic effects on the output of the basal ganglia. We optogenetically stimulated the direct and indirect pathways in mice and measured their influence on perceptual decisions and neuronal activity in the cortex. In a task in which mice had to detect a visual stimulus, unilateral direct-pathway stimulation increased the probability of lick responses to the non-stimulated side and increased cortical activity globally. In contrast, indirect-pathway stimulation increased the probability of licks to the stimulated side and decreased activity in visual cortical areas. To probe the possible role of the two pathways in working memory, we trained the mice to report the location of a stimulus with licking one of two spouts, after a memory delay. Direct-pathway stimulation prior to and during the memory delay enhanced both the neural representation of a contralateral visual stimulus and the number of contraversive choices, whereas indirect-pathway stimulation had the opposite effects, in accordance with an antagonistic influence of the direct and indirect pathways on licking direction. Our results demonstrate how these two pathways influence perceptual decisions and working memories, and modify activity in the cerebral cortex.

**One sentence summary:** Visuomotor transformations are influenced antagonistically by the direct and indirect pathways of the basal ganglia during visual detection and working memory tasks

## Introduction

We constantly make decisions based on sensory information. Sometimes a stimulus requires an immediate response, and in other situations responses can be postponed to a later point in time, so that information must be kept “online” in working memory. Previous studies in non-human primates demonstrated that working memories are associated with delay activity in multiple brain regions, reflecting the properties of previously presented sensory stimuli (Curtis and D’Esposito, 2003; Fuster and Alexander, 1971, 1973; Fuster and Jervey, 1981; Van Kerkoerle et al., 2017; Kubota and Niki, 1971; Mendoza-Halliday et al., 2014; van Vugt et al., 2020). These results were replicated in the mouse, where visual, multisensory and frontal brain regions maintain activity related to previously presented stimuli (Goard et al., 2016; Guo et al., 2017). Importantly, persistent activity does not only reflect the memory of previous stimuli, but it also plays a role in the planning of future behavior, decision making, and in the retrieval of associations between related concepts (Baddeley, 1992; Christophel et al., 2017; Ding and Gold, 2010; Osaka et al., 2012; Peixoto et al., 2021; Rainer et al., 1999).

The finding that persistent activity occurs in many brain regions suggests that it may be an emergent property of distributed brain networks (Christophel et al., 2017; Dotson et al., 2018; Voitov and Mrsic-flogel, 2022). Interestingly, brain regions that are of importance in tasks that rely on working memory are often also involved in simpler decision-making tasks. Example brain regions in mice involved in working memory and simple decisions include region ALM of the frontal cortex (Chen et al., 2017; Esmaeili et al., 2021; Goard et al., 2016), the dorsomedial striatum (Wang and Krauzlis, 2020; Wang et al., 2018, 2021) and the lateral striatum (Lee et al., 2020). However, it remains unclear how different brain regions orchestrate their activity to maintain working memories. The present study examines the possible role of the basal ganglia as a coordinator of persistent activity for working memory.

The cortex, basal ganglia and thalamus are organized in an anatomical loop. Different cortical regions project to distinct regions of the striatum, which is the primary input structure of the basal ganglia. This segregation is maintained throughout successive basal-ganglia nuclei (Alexander et al., 1986; Haber, 2016; Lee et al., 2020; McGeorge and Faull, 1989; Saint-Cyr et al., 1990), which ultimately project to the thalamus that, in turn, loops the information back to cortical regions that provided the input (Foster et al., 2021). In line with this connectivity pattern, activity in different striatal domains correlates with activity in specific regions of the cortex (Choi et al., 2012; Peters et al., 2021). The activity of neurons in dorsomedial regions of the striatum correlates with activity in visual areas and midline frontal regions, whereas the activity of neurons in dorsolateral regions of the striatum correlate with activity in motor regions, such as the anterior lateral motor cortex (ALM). Previous studies suggested that the loops from the cortex through the basal ganglia could play a role in the maintenance of persistent activity for working memory (Wang, 2001; Wang et al., 2021). In accordance with this view, the nuclei of the basal ganglia are connected with frontal cortical regions that exhibit persistent activity during working-memory tasks (Saunders et al., 2015) and striatum-projecting neurons in the medial prefrontal cortex contribute to the maintenance of spatial working memories (Chernysheva et al., 2021). Furthermore, neurons of the basal ganglia themselves exhibit persistent activity during working memory tasks (Hikosaka and Wurtz, 1983; Hikosaka et al., 1989; Mushiake and Strick, 1995).

The cerebral cortex projects to GABAergic neurons in the striatum that belong to the direct and indirect pathways (**Figure 1B**), and express different dopamine receptors. Direct-pathway striatal projection neurons (dSPNs) express D1 dopamine receptors, whereas indirect-pathway projection neurons (iSPNs) express D2 dopamine receptors (see **Table S1** for abbreviations). The dSPNs and iSPNs have opposite effects on activity in basal-ganglia output structures, leading to different influences on action initiation and execution (Nonomura et al., 2018; Tecuapetla et al., 2016). In rodents, dSPNs inhibit neurons in the substantia nigra pars reticulata and the entopeduncular nucleus (Smith et al., 1998) (**Figure 1B**), which, in turn, inhibit nuclei of the brainstem, the superior colliculus and the thalamus (Wang et al., 2021). Hence, the direct pathway has two inhibitory synapses so that dSPNs have an overall excitatory effect on neurons in the superior colliculus, thalamus and cortex. The iSPNs inhibit neurons in the globus pallidus, which inhibits the substantia nigra pars reticulata and entopeduncular nucleus (Smith et al., 1998). Hence, there is a third GABAergic connection in the indirect pathway so that the net effect of iSPNs on the superior colliculus, thalamus and cortex is inhibitory (Lee and Sabatini, 2021; Lee et al., 2020), which can explain their possible role in the inhibition of actions (Cruz et al., 2022). Besides receiving input from all cortical areas, the striatum also receives strong dopaminergic inputs, which influence synaptic plasticity, allowing the basal ganglia circuits to adapt during reinforcement learning (Cox and Witten, 2019; Schultz, 2016; Schultz et al., 1997). Indeed, the inputs from the cortex to the striatum are potentiated when new tasks are learned (Peters et al., 2021; Xiong et al., 2015; Znamenskiy and Zador, 2013).

**Figure 1.**
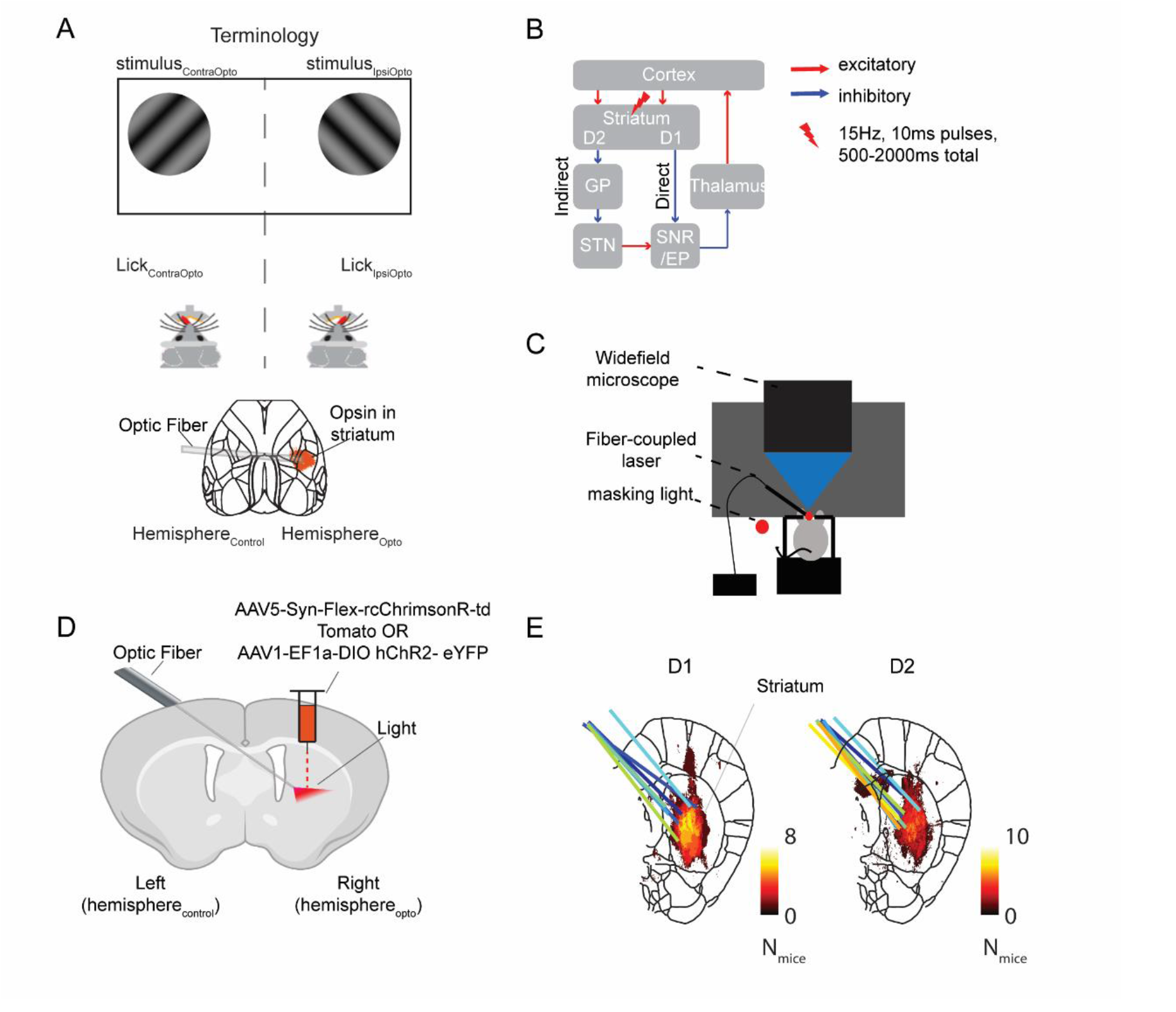
Optogenetic stimulation of the striatum during widefield imaging of the cortex. A) We optogenetically stimulated the striatum of right hemisphere (hemisphere_Opto_). The view of the cortex of the non-stimulated hemisphere_Control_ was partly occluded by the ferrule holding the optic fiber and the cement to hold it in place. Visual stimuli were presented in the hemifield contralateral (stimulus_ContraOpto_) or ipsilateral (stimulus_IpsiOpto_) to hemisphere_Opto_. The mouse licked a spout in a contraversive or ipsiversive direction relative to hemisphere_Opto_. B) Schematic of the loop from cortex to basal ganglia, then to the thalamus and back to cortex. We expressed an excitatory opsin in the direct (dSPN neurons in D1-cre mice) or indirect (iSPN neurons in D2-cre mice) pathway. C) Set-up for widefield calcium imaging during optogenetic stimulation. Mice were head-fixed under the widefield microscope and a fiber coupled to a laser was aimed at the striatum. A masking light of approximately the same wavelength blinked at the same frequency and with the same pulse width as the laser, but at random time points. D) A viral vector was injected in the striatum of hemisphere_Opto_ of D1 or D2-cre mice. The tip of the optic fiber ended above the region with virus expression (image created with BioRenderer). E) Expression of the excitatory opsin in the striatum (coronal slice +0.14mm anterior to Bregma) across mice for D1 (left) and D2 (right) mice. Histology from individual mice was aligned to the Allen Brain atlas. The colour depicts the number of mice for which a certain pixel showed expression of the excitatory opsin above a threshold (see Methods). The coloured lines show the tracts of the optic fiber in individual mice (for expression in individual mice see Figure S1).

The position of the basal ganglia enables them to coordinate decision making. They receive sensory input from the cortex, reward and motivation signals through dopaminergic inputs, and they can gate whether activity is fed back to the cortex via the thalamo-cortical loop by balancing inputs via the direct and indirect pathways. Furthermore, computational studies suggested that the basal ganglia could control the contents of working memory by switching the two complementary pathways on and off (Frank et al., 2001; Wise et al., 1996). It is therefore of great interest to better understand the role of the basal ganglia in working memory.

In this study, we asked how the basal ganglia influence persistent activity in the cortex related to perceptual decisions and working memory. We injected a virus unilaterally to express an excitatory opsin either in the direct or indirect pathway and measured behavioral responses and neuronal activity across the dorsal cortex in adult mice using widefield imaging and optogenetics. To understand the role of this circuitry in different behavioral states, we tested the influence of the two pathways during a task requiring detection of a visual stimulus and a task in which mice had to memorize a stimulus during a delay.

## Results

We set out to identify the roles of the direct and indirect pathways in visuomotor transformations with and without a memory requirement (**Figure 1A**). In the first experiment, we tested the influence of direct and indirect pathway activation on activity of the cerebral cortex outside a task. In the subsequent experiment, we investigated how activity of these pathways influences behavioral responses and cortical activity in a visual detection task. The last experiment examined persistent activity in the cortex during a working memory task, and how it is influenced by the direct and indirect pathways.

We included 6 Thy1-5.17 GCaMP6f mice (Dana et al., 2014), 8 D1-cre, 10 D2-cre, 9 Thy1-5.17 GCaMP6f X D1-cre, and 7 Thy1-5.17 GCaMP6f X D2-cre of both sexes, aged 2-6 months at the start of the experiment (**Table S2**). GCaMP6f is a calcium sensor for widefield imaging, which in Thy1 mice is globally expressed in excitatory neurons (Dana et al., 2014; Ren and Komiyama, 2021). We used a method that made the skull transparent (clear-skull technique, Guo et al., 2014b) so that we could image neuronal activity of the dorsal cortex by measuring the change in GCaMP6f fluorescence from baseline activity (ΔF/F).

To activate the direct or indirect pathway, we injected D1-cre x Thy1-GCaMP6f (D1) and D2-cre x Thy1-GCaMP6f (D2) mice with AAV5-syn-Flex-rcChrimsonR-tdTomato or AAV1-EF1a-DIO-hChR2-eYFP in the ventromedial part of the dorsal striatum of the right hemisphere (hemisphere_Opto_, **Figure 1**). The virus induced the expression of excitatory opsins Chrimson or ChR2. These proteins cause depolarization of neurons upon stimulation with light. The expression of the opsin was conditional on Cre activity, so that it occurred only in dSPNs in D1 mice and in iSPNs in D2 mice. We optogenetically stimulated the striatum at 15Hz with 10ms light pulses. We pooled the results across the two opsins as they were similar. To direct light to the striatum of hemisphere_Opto_, we implanted an optic fiber at an angle, entering the skull in the non-injected hemisphere (hemisphere_Control_). This allowed us to image the dorsal cortex of hemisphere_Opto_ entirely, whereas the view of a central region of hemisphere_Control_ was obstructed by the implant (**Figure 1C,D**). Upon completion of the experiments, we examined the expression of the opsin and the location of the optic-fiber tips (**Figure 1E**, **Figure S1**). We only analyzed the data of D1- and D2-mice in which the opsin was expressed in the striatum and the fiber correctly illuminated this brain region. Control mice underwent the same surgery and experimental procedures, but histology revealed no virus expression in or near the striatum.

### The influence of direct and indirect pathway stimulation on cortical activity

To examine how the direct and indirect pathway modulate cortical activity outside the context of a task, we optogenetically stimulated the striatum for a total duration of 1 second while imaging activity in the dorsal cortex, i.e. the regions of the cortex that could be imaged through the cleared skull. The mice were head-fixed in the setup but did not perform a task and did not have access to a lick spout. In D1-mice (N=5), optogenetic stimulation of dSPNs caused a significant increase in activity (ΔF/F of the calcium signal) of large regions of hemisphere_Opto_ and hemisphere_Control_ (**Figure 2A,B**) (mixed effects model of stimulation induced change in ΔF/F with factors cortical area and hemisphere, and mouse as random factor; intercept: F_1,7930_=11.3, p<0.001). The most notable differences in the level of stimulation induced activity between areas were in the lateral visual areas, which were strongly activated in hemisphere_Opto_ but not in hemisphere_Control_ (interaction between area and hemisphere: F_9,7930_=2.18, p<0.05; post-hoc Wald tests of coefficients, p<0.05, **Figure 2C**). In D2-mice (N=3) we did not observe a consistent change in ΔF/F activity upon iSPN stimulation outside the context of a task (mixed effects model, p>0.05).

**Figure 2.**
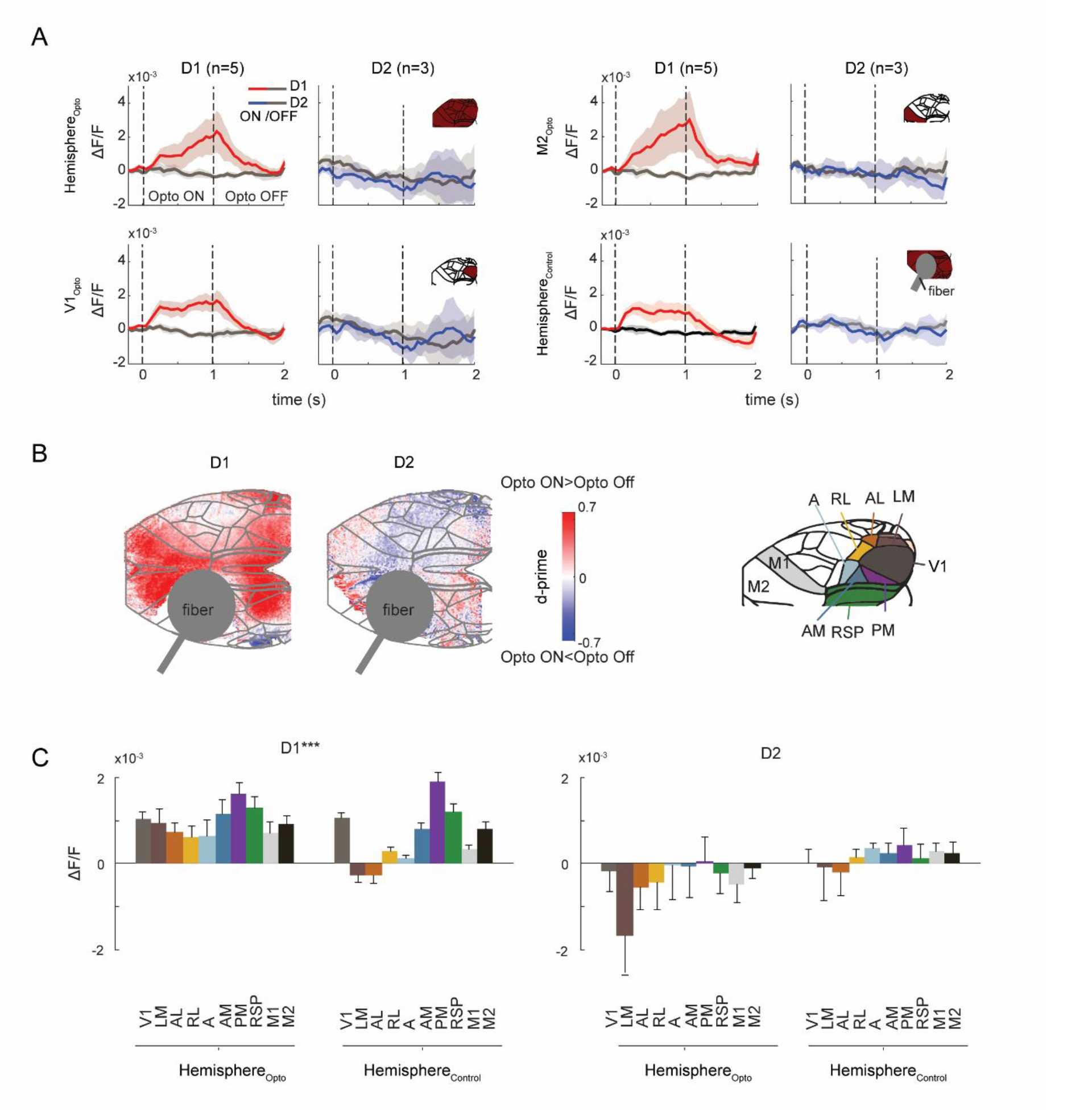
Effect of direct and indirect pathway stimulation on cortical activity. A) Average time-course of the GCaMP signal for the entire hemisphere_Opto_ (top row), area V1 (second row), area M2 (third row) and the entire hemisphere_control_ (bottom row) in D1 (left panels) and D2 (right panels) mice. Grey traces show ongoing activity and red/blue traces the activity induced by optogenetic stimulation. Shaded regions denote s.e.m. B) The effect of stimulation was quantified with d-prime in D1 (left panel) and D2 (right panel) mice. Red colors indicate cortical excitation and blue colors cortical inhibition. C) ΔF/F in D1 (left) and D2 (right) mice during optogenetic stimulation in different areas (colors are indicated in the schematic brain inset). Mixed-effects model with factors cortical area and hemisphere revealed a significant increase in activity for D1 mice (intercept: F_1,7930_=11.30, p<0.001) and a significant interaction between area and hemisphere (F_9,7930_=2.18, p<0.05). Post-hoc Wald tests revealed that optogenetic stimulation of dSPNs caused a larger increase of activity in V1 than in lateral visual areas (LM, AL and RL) and motor regions (p<0.05), but smaller than activity in medial visual areas (PM and AM) (p<0.05). These differences between areas were more pronounced in hemisphere_control_.

### The direct and indirect pathways influence lick responses in a visual detection task

Next, we tested if the direct and indirect pathways play a causal role in the mapping of visual stimuli onto motor responses using a visual-detection task in eight D1- and eight D2-mice. A visual stimulus appeared either in the hemifield contralateral to the stimulated striatum (stimulus_ContraOpto_) or in the hemifield on the ipsilateral side (stimulus_IpsiOpto_). The mice could lick either side of a two-sided lick spout to obtain a reward (**Figure 3A**). The reward was given upon the first contact of the tongue with the spout after stimulus onset, and the water came from the same side as the visual stimulus, even when the first lick was on the other side. In 20% of the trials, optogenetic stimulation started 0.5s prior to visual stimulus onset and lasted for 2 seconds (see Materials and Methods) and in another 20% of trials it started immediately after the first lick.

**Figure 3.**
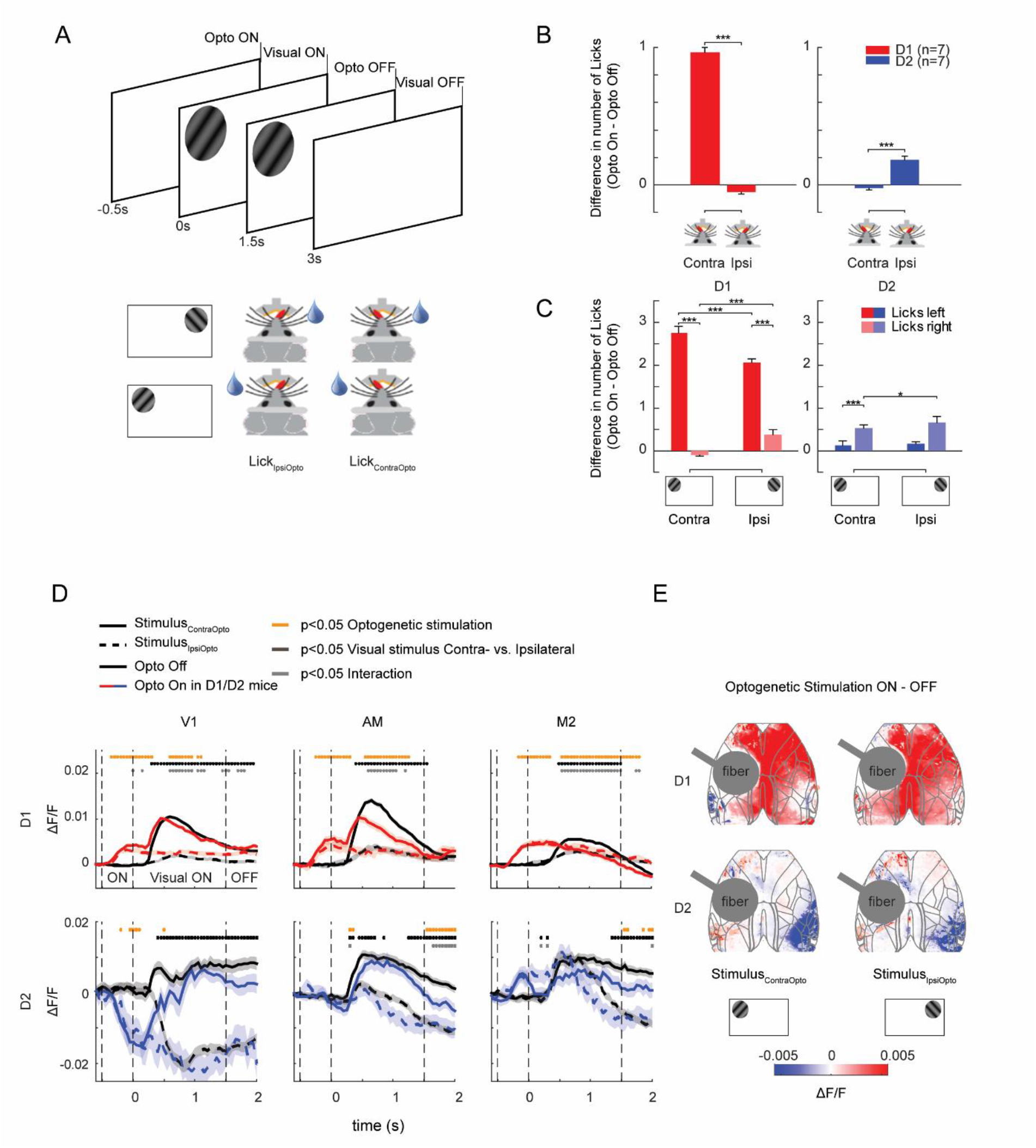
Effect of optogenetic stimulation of the striatum in the visual detection task. A) Visual detection task. Mice saw a drifting grating in the hemifield contralateral (stimulus_ContraOpto_) or ipsilateral (stimulus_IpsiOpto_) to the stimulated striatum (hemisphere_Opto_). Optogenetic stimulation occurred on 40% of the trials and started in the baseline period (i.e. -0.5 to 0s relative to the visual stimulus onset) on half of these trials, and immediately after the first lick on the other half. We compared trials without optogenetic stimulation (Opto Off) to those with optogenetic stimulation in the baseline period (Opto On). Lick responses to either lick spout were rewarded on the same side as the visual stimulus. B) Influence of optogenetic stimulation on the mean number of licks in the baseline period for D1 (red) and D2 (blue) mice. Significance was determined with a mixed effects model and post-hoc Wald tests of coefficients on the interaction of optogenetic stimulation and lick side; corrected for multiple comparisons. Error bars denote s.e.m. of the optogenetic stimulation effect. C) Same as B, but during the presentation of stimulus_ContraOpto_ and stimulus_IpsiOpto_. Bars with darker (lighter) shade represent contraversive (ipsiversive) licks. Error bars denote s.e.m. of the difference in lick numer between optogenetic stimulation on and off. D) Time-courses of GCaMP signal of three example areas in hemisphere_Opto_ in D1 (N=4) and D2 mice (N=3). Solid (dashed) traces, activity induced by stimulus_ContraOpto_ (stimulus_IpsiOpto_). The different epochs (optogenetic stimulation on, visual stimulus on, both off) are indicated by vertical dashed lines. Shaded area denotes s.e.m. Orange dots, main effect of optogenetic stimulation (p<0.05; mixed effects model). Black dots, main effect of the location of the visual stimulus (p<0.05). Grey dots, interaction between these factors (p<0.05). E) Average effect of optogenetic stimulation on cortical activity (time window 0-1500ms) during stimulus_ContraOpto_ (left column) and stimulus_IpsiOpto_ (right column). In all panels *, p<0.05; ***, p<0.001.

Our analysis focused on the trials in which stimulation started before stimulus onset (the results on trials in which stimulation started after the first lick were similar, see Supplementary Analysis). For statistical analysis, we used a mixed effects model predicting the number of licks assuming a Poisson distribution, with factors optogenetic stimulation and lick side, and mouse as a random effect. In D1 mice, optogenetic stimulation increased the number of contraversive licks in the pre-stimulus epoch (interaction between lick side and optogenetic stimulation: F_1,7962_=307, p<0.001; main effect of lick side: F_1,7962_=92, p<0.001, **Figure 3B**, **Figure S2A**). In contrast, optogenetic stimulation in D2 mice before stimulus appearance led to a larger increase in the number of ipsiversive licks than contraversive licks (interaction optogenetic stimulation and lick side: F_1,6956_=36, p<0.001; main effect of lick side: F_1,6956_=97, p<0.001).

Does stimulation of the direct and indirect pathway induce a lateralized licking response, or does it interact with the location of the visual stimulus? To examine this, we analyzed the effect of optogenetic stimulation on licking in the epoch after the stimulus had appeared. In D1 mice, optogenetic stimulation indeed interacted with the position of the visual stimulus (mixed effects model with stimulus location as additional factor, main effect optogenetics: F_1,5834_=884, p<0.001; interaction with visual stimulus: F_1,5834_=84, p<0.001, and interaction with lick side: F_1,5834_=114, p<0.001). dSPN activation increased contraversive licks for both visual stimuli and in case of stimulus_IpsiOpto_ there was also a small increase in the number of ipsiversive licks (post-hoc Wald tests of coefficients for the difference between stimuli, p<0.001; **Figure 3C**). A similar, albeit weaker interaction occurred in D2 mice (main effect of optogenetics: F_1,5220_=5.7, p<0.05; three-way interaction between optogenetic stimulation, visual stimulus, and lick side: F_1,5220_=14.6, p<0.001). Specifically, with iSPN stimulation mice made more ipsiversive licks when an ipsilateral stimulus was presented compared to when a contralateral stimulus was presented (post-hoc Wald tests of coefficients p<0.05; **Figure 3C**). This interaction with the stimulus position implies that optogenetic stimulation did not simply force a lateralized lick response. Furthermore, optogenetic stimulation did not impair the ability of mice to switch licking to the spout that delivered the water when the first lick was toward the other one (see Supplementary Analysis).

Taken together, these results demonstrate that the direct and indirect pathways antagonistically influence the direction of lateralized lick-responses, in a manner that depends on the visual input and the location of the reward. We next examined how the activation of dSPN and iSPN neurons influences cortical activity.

### Influence of striatal pathways on cortical activity during the visual detection task

We imaged four D1 mice and three D2 mice to examine how stimulation of the direct and indirect pathways influences cortical activity during the task. Optogenetic stimulation of dSPNs caused a pronounced and global increase in cortical activity (**Figure 3D,E**; **Figure S2C**). **Figure 3D** illustrates dSPN activation in example cortical areas (compare red and black traces; orange, black and grey symbols show the results of a mixed effects model per time point with ∼250 trials per mouse, ps<0.05). After the initial increase in cortical activity before the visual stimulus, V1 activity became similar with and without dSPN stimulation, whereas activity in AM and M2 even dropped below the level without dSPN activation (**Figure 3D**), causing an interaction between optogenetic stimulation and the position of the visual stimulus (grey symbols in **Figure 3D**).

We next examined the influence of optogenetic stimulation of iSPNs on cortical activity. Prior to stimulus onset, iSPN activation decreased V1 activity in hemisphere_Opto_, and increased activity in several cortical regions of hemisphere_Control_ (p<0.05 for time points -0.5 to 0.5s) (**Figure 3D,E; Figure S2C**). At later time points, optogenetic stimulation caused a more general decrease in activity that was most pronounced in the visual cortex (p<0.05 for the main effect of optogenetic stimulation). Stimulation of iSPN neurons increased activity in many areas of hemisphere_Control_ before stimulus onset but tended to decrease activity thereafter (**Figure 3E; Figure S2C**). These results, taken together, demonstrate that direct pathway stimulation causes a relative widespread increase of cortical activity and that indirect pathway stimulation decreases activity of visual cortical areas.

### Training changes the dSPN stimulation effect

Does training influence the effect of direct pathway stimulation on cortical activity outside the task? To examine the effect of training, we compared cortical activity elicited by dSPN stimulation before and after training (N=2 mice). Training boosted dSPN excitation (two-way repeated measures ANOVAs with factors area and training history, main effect of training history: F_1,1480_=8.8, p<0.01 for mouse A; F_1,980_=33, p<0.001 for mouse B, **Figure S3A,B**). Strikingly, pixels that were activated by the visual detection task were also activated most by dSPN stimulation after training (**Figure S3B**), an effect that was validated with a correlation analysis (Supplementary Analysis).

### Persistent neuronal activity during a working memory task

Our last experiment explored the influence of dSPNs and iSPNs on a task that required working memory, perceptual decisions, and motor responses. We used a two-alternative-forced-choice task with a delay between the visual stimulus and the motor response (**Figure 4A**). The visual stimulus (same as described above) was presented on the left or right. After a memory delay of 1.5s, the mice had to report the location of the visual stimulus by licking the corresponding side of the two-sided lick spout. In this version of the task, a reward was only given if the first lick was on the same side as the visual stimulus. Most of the mice (77%) performed the task with a 1.5s delay above chance level after 40-100 training sessions (5 training sessions per week), with 150-250 trials per session (**Figure 4B,C**; p<0.001 for both visual stimuli). Mice that did not learn the task were excluded from further analysis (10 mice remained).

**Figure 4.**
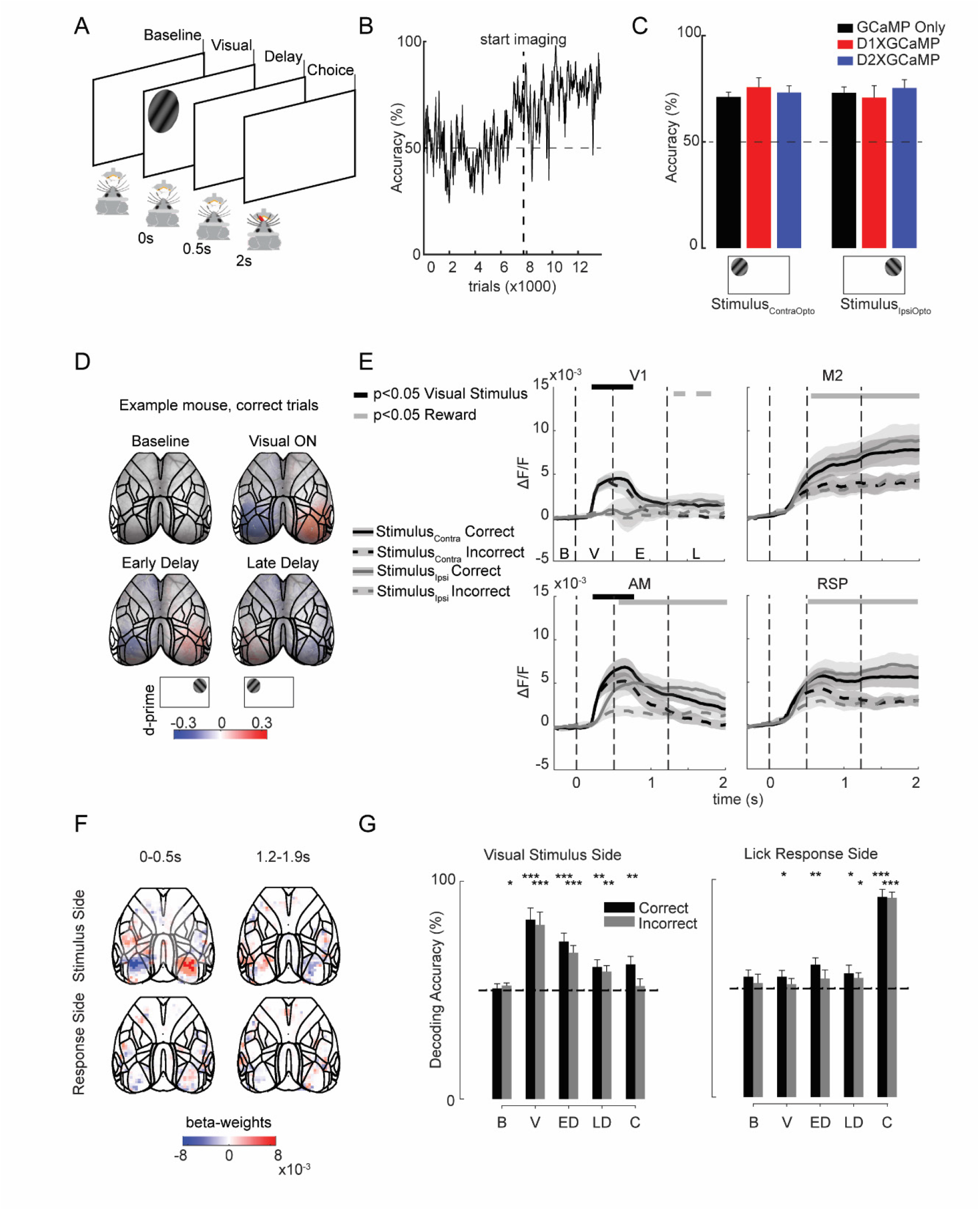
Cortical activity in the working memory task. A) Working memory task. We presented a drifting grating in the left or right hemifield for 500ms. After a delay of 1500ms, a two-sided spout was placed close to the mouth of the animal so that they could lick. Licks on same side as the stimulus were rewarded. B) Learning curve of one of the mice. C) Average accuracy (black: GCaMP only, red: D1 x GCaMP, blue: D2 x GCaMP). D) D-prime for the side of the stimulus in successive time windows. Colored regions denote significant d-prime values (t-test, p<0.05, uncorrected). E) Time-course of GCaMP fluorescence in four example cortical areas. Solid (dashed) traces, correct (erroneous) trials. Black (grey) traces show activity elicited by a stimulus in the contralateral (ipsilateral) hemifield. Letters on the x-axis refer to the different (baseline, visual stimulus presentation, early and late delay). Black lines above the graph show significant main effects of stimulus side, grey lines a main effect of correct vs. error trials in a two-way ANOVA per time point. Note the higher activity in M2 and RSP in correct trials. F) Beta-weights of the MALSAR decoder for side of stimulus (upper panels) and upcoming lick direction (lower panels) in an early and later time window (0-0.5s and 1.2-1.9s after visual stimulus onset). Red pixels are positively correlated with the stimulus in the left hemifield (stimulus_ContraOpto_) or left lick (lick_ContraOpto_) and blue pixels with the opposite stimulus and lick. Fully opaque pixels show beta-values significantly different from zero across mice (N=10, t-test, p<0.05, uncorrected). G) Cross-validated decoding accuracy (mean +/- s.e.m.) across mice (N=10) for side of the visual stimulus (left panel) and lick response (right panel). Black bars for correct, and grey bars for incorrect trials. Decoding accuracy for stimulus and lick response side depended on the time-window (p<0.05; MANOVA), but not on the trial outcome (correct/error). *, p<0.05; **, p<0.01;***, p<0.001 in post-hoc t-tests.

The task activated regions of the visual and motor cortex, in accordance with previous work (Chen et al., 2017; Goard et al., 2016; Orsolic et al., 2021). We analyzed the results with a repeated measures ANOVA for every time point, with factors correct/error and visual stimulus position across hemispheres (N=15 hemispheres). As expected, the visual stimulus initially drove activity in the contralateral visual cortex (p<0.05; **Figure 4D,E**). In the delay epoch, we observed persistent activity in many cortical areas. To examine if the persistent activity was related to working memory, we compared correct and erroneous trials (**Figure 4E**). Persistent activity was higher on correct trials, in particular in the higher cortical areas (ps<0.05 for multiple time points; compare dashed traces to continuous traces in **Figure 4E**), in accordance with a role in working memory.

Next, we examined how delay activity encoded the position of the visual stimulus. In the early phase of the delay epoch, activity was stronger in visual areas contralateral to the visual stimulus (p<0.05 for multiple time points early in the delay, N=15 hemispheres; **Figure 4D,E**). Interestingly, in a later phase of the delay the pattern inverted so that there was a non-significant trend for activity elicited by the memory of an ipsilateral visual stimulus to be stronger (ps>0.05; repeated measures ANOVA for every time point). This inversion may represent a form of adaptation, but it may also relate to memory encoding.

We used a multi-output decoding approach (MALSAR) to decode stimulus position and the upcoming licking direction (Zhou et al., 2012). As expected, the decoding weights in visual areas inverted in the late delay epoch (**Figure 4F**). The accuracy of decoding varied across trial time (**Figure 4G**) (N=10 mice, ANOVA with factors correctness and time window, for stimulus position decoding: F_4,90_=22.1, p<0.001 and for lick direction: F_4,90_=56.3, p<0.001). Decoding of the stimulus was best while it was visible and the decoding accuracy decreased later in the trial, although it continued into the delay epoch (**Figure 4G**, left panel, p<0.01 post-hoc t-tests). The accuracy of decoding the upcoming lick direction gradually increased during the trial (**Figure 4G**, right panel, p<0.05 post-hoc t-tests). These results, taken together, indicate that persistent activity in the cortex carries information about the previously presented stimulus as well as the upcoming lick response.

### The direct and indirect pathways influence accuracy during a working memory task

We tested the influence of optogenetic stimulation on accuracy in two D1-mice and three D2-mice in the working memory task. Without optogenetic stimulation, the accuracy of these mice was comparable to that of the control mice (**Figure 4C**). We applied optogenetic stimulation in one of the following epochs: just before the visual stimulus (-0.5-0s), during the visual stimulus (0-0.5s), the delay (1-1.5s) or the response epoch (2-2.5s).

The influence of optogenetic dSPN stimulation depended on the position of the visual stimulus and the epoch (mixed effects model, significant interaction between epoch and stimulus side: F_5,12_=10.9, p<0.001). Specifically, stimulation of dSPNs during the visual stimulus presentation and delay epochs increased contraversive choices, boosting the accuracy for stimulus_ContraOpto_ and decreasing it for stimulus_IpsiOpto_ (**Figure 5A**). Interestingly, this effect also occurred when we applied optogenetic stimulation just prior to the visual stimulus. Stimulation of iSPNs in D2-mice had the opposite effect by increasing the number of ipsiversive choices (mixed effects model as above, significant interaction: F_5,24_=9.6, p<0.001). It increased the accuracy for stimulus_IpsiOpto_ and decreased it for stimulus_ContraOpto_ in the pre-stimulus epoch, during the visual stimulus and during the delay. Stimulation of dSPNs and iSPNs did not influence the number of omission trials without a lick response (**Figure 5A**) (mixed effects model, no significant effects: all ps>0.05). Hence, activation of the direct and indirect pathways prior to the late delay had a pronounced influence on the decision that was taken at the end of the trial.

**Figure 5.**
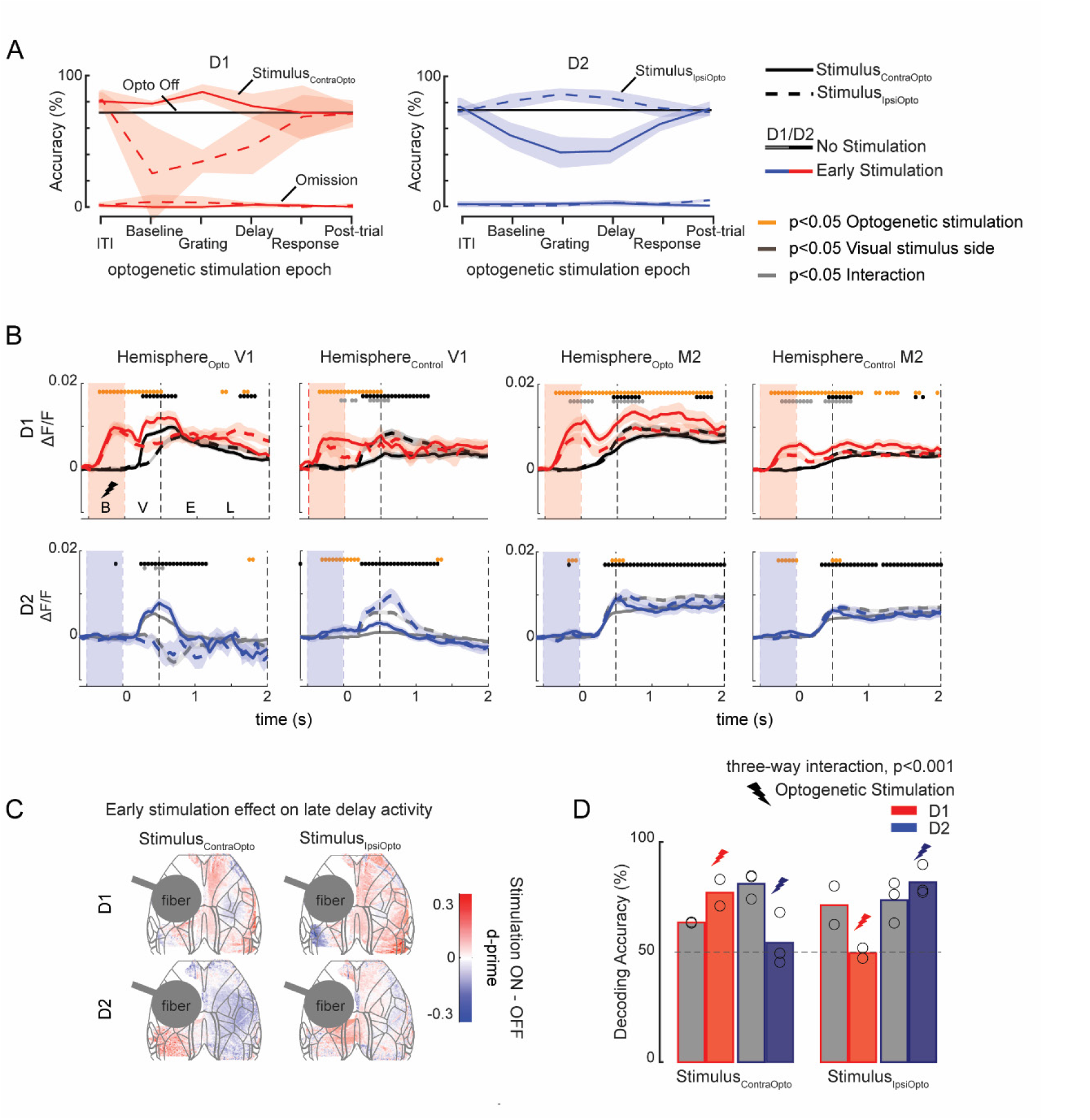
Optogenetic stimulation of the striatum in the working memory task. A) Influence of optogenetic stimulation in different time-windows (x-axis) on the accuracy of D1 (left panel) and D2 (right panel) mice. The black horizontal line shows accuracy in trials without optogenetic stimulation. Blue and red solid (dashed) lines show accuracy (top lines) and percentage omission (bottom lines) for stimulus_ContraOpto_ (stimulus_IpsiOpto_). Optogenetic stimulation did not alter the percentage of omissions. Accuracy increased (decreased) for stimulus_ContraOpto_ (stimulus_IpsiOpto_) in D1 mice, and the opposite was true for D2 mice. B) Influence of optogenetic stimulation (colored traces) in the baseline period (from -500-0ms) on M2 and V1 activity. Traces show average ΔF/F and shaded regions s.e.m. Solid (dashed) lines represent trials with stimulus_ContraOpto_ (stimulus_IpsiOpto_). Mixed-effects models per time point showed significant main effects of optogenetic stimulation (orange circles, p<0.05), visual stimulus side (black circles, p<0.05) and an interaction between these factors (grey circles, p<0.05) (see Figure S3 for optogenetic stimulation during the visual stimulus presentation). C) D-prime of the optogenetic effect (pooled across all early optogenetic stimulation trials) during the late delay (1200-1900ms after stimulus onset). D) We trained a model to decode stimulus position based on cortical activity during the late delay on trials without optogenetic stimulation. We then tested the model on late delay cortical activity of trials with early optogenetic stimulation (red/blue). We also included a separate test-set of trials without stimulation (gray). Depicted is the percentage of trials that the model predicted the memory representation of stimulus_ContraOpto_ (left) and stimulus_IpsiOpto_ (right) correctly. The increase in accuracy for stimulus_ContraOpto_ for D1 mice (bars with red outline) after early optogenetic stimulation of the striatum shows that cortical activity in the late delay is biased to represent stimulus_ContraOpto_, and vice versa for D2 mice (bars with blue outline). A three-way ANOVA revealed a significant interaction between optogenetic stimulation, visual stimulus side and genotype (p<0.001) (see Figure S5 for more details). Circles represent decoding accuracy for individual mice.

### The direct and indirect pathways influence neuronal representations of working memory

To analyze the influence of direct and indirect pathway stimulation on cortical activity during the working memory task, we combined all trials in which it strongly influenced behavior. Specifically, we included trials in which the optogenetic stimulation was given in the (pre-stimulus) baseline epoch (**Figure 5B**), during the presentation of the visual stimulus (**Figure S4A**) and early in the delay.

dSPN stimulation caused an immediate increase in cortical activity (see **Figure 5B** for two example areas, V1 and M2). Interestingly, optogenetic stimulation early in the trial had an effect on cortical activity during the late delay [1.5-1.9s] in both D1 and D2 mice (mixed effects model performed per time point: main effect of optogenetic stimulation as orange symbols and interaction with visual stimulus position as gray symbols, p<0.05 in **Figure 5B, Figure S4**). Activation of dSPNs increased the activity in M2 in both hemisphere_Opto_ and hemisphere_Control_, although the increase in hemisphere_Opto_ was more pronounced. Activation of iSPNs stimulation initially increased the visually driven response in visual cortex, but during the late delay we observed a decrease in persistent activity for stimulus_ContraOpto_ in hemisphere_opto_ (**Figure 5B,C, Figure S4**). The influence on the decision could have altered the memory of the stimulus, the representation of the upcoming lick response, or both.

To examine the influence of the striatal pathways on the working memory representation, we trained decoding models using cortical activity from trials without optogenetic stimulation. We then tested how optogenetic stimulation influenced decoding during the late delay. To gather sufficient statistical power, we pooled together trials with optogenetic stimulation in the baseline epoch, during the visual stimulus and early in the memory delay. Without optogenetic stimulation, decoding accuracy for the location of the visual stimulus was significant in all mice (grey bars in **Figure 5D**). Early dSPN stimulation biased the decoder to output stimulus_ContraOpto_ more often during the late delay. Early iSPN stimulation biased the decoder oppositely to output stimulus_IpsiOpto_ (ANOVA, significant interaction of the three factors optogenetic stimulation, visual stimulus side and genotype: F_1,12_ =20.2, p<0.001, **Figure 5D**, see **Figure S5B** for data of individual mice). This result indicates that optogenetic stimulation of the direct and indirect pathways influenced the memory of the visual stimulus.

Optogenetic stimulation did not influence decoding of lick direction, even though the number of contralateral licks increased with dSPN stimulation and decreased with iSPN stimulation (**Figure 5A**). It seems likely that this is caused by the correct decoding of altered lick responses, without a strong influence of optogenetic stimulation on the relation between licking direction and wide-field activity.

## Discussion

We investigated the influence of direct and indirect pathways of the basal ganglia on visuomotor transformations and activity in the cerebral cortex. In the absence of a task, dSPN stimulation led to a global increase of cortical activity, whereas stimulation of iSPNs only had a weak and more local effect. In the visual detection task, stimulation of dSPNs increased the number of contraversive licks (**Figure 3**). It also caused an overall increase in cortical activity, with an extra enhancement of activity in area M2 if the visual stimulus fell in the hemifield contralateral to hemisphere_Opto_. In contrast, iSPN stimulation increased the number of ipsiversive licks and it decreased neuronal activity in the visual cortex of hemisphere_Opto_. Training in the visual detection task enhanced the influence of optogenetic stimulation on cortical activity, with an increased resemblance to the activity pattern that occurred during the task. In our final experiment, we tested the influence of dSPNs and iSPNs in a working memory task. We first established signatures of cortical activity that were associated with the visual stimulus, its memory, and the direction of the lick response. We then applied optogenetic stimulation to the striatum and found that direct-pathway activation increased the number of future contraversive choices. This effect was observed with optogenetic stimulation in the epochs just before the stimulus appeared, during the presentation of the visual stimulus, and during the delay between stimulus and lick response, but not in epochs seconds before the trial or during the lick response itself. When we examined cortical activity at the end of the delay period, we found that dSPNs stimulation during earlier time points caused activity to resemble a working memory of a visual stimulus presented contralateral to hemisphere_Opto_, in accordance with the increase in contraversive choices. In contrast, iSPN activation in the working memory task increased future ipsiversive choices, an effect that was associated with a cortical activity pattern resembling the working memory of a visual stimulus ipsilateral to hemisphere_Opto_.

Previous studies have linked the basal ganglia to the execution of motor sequences and the mapping of stimuli onto responses (Graybiel, 1998; Groenewegen, 2003) through reinforcement learning (White, 1997). Indeed, the influence of the dorsal striatum on behaviour depends on the task and the actions that are associated with a reward (Lee and Sabatini, 2021; Lee et al., 2021; Wang et al., 2018). We found that the direct and indirect pathway in the striatal region that we targeted (the ventromedial part of the dorsal striatum) promote contraversive and ipsiversive licks, respectively. The effect of dSPN and iSPN activation depended on the location of the visual stimulus (see also Bolkan et al., 2022). We therefore attribute these influences to the mapping of visual stimuli onto motor responses. Hence, this striatal region does not instruct lick responses at the motor level and, accordingly, the mice could switch to the other direction if the first lick was in the wrong direction.

Our finding in the no-delay task support Lee et al. (2020), who trained mice to report one of two auditory tones with a left or right lick response. In their experiment, dSPN stimulation also increased contraversive licks, which was associated with more activity in the region of the motor cortex related to licking responses. They recently demonstrated that iSPNs influence the superior colliculus (Lee and Sabatini, 2021), which also influences licking (Rossi et al., 2016). The iSPNs inhibit the ipsilateral and excite the contralateral superior colliculus, thereby increasing ipsiversive licks. Similarly, dSPNs excite the motor cortex, whereas iSPNs have an inhibitory influence (Oldenburg and Sabatini, 2015).

Our results go beyond these previous studies by demonstrating an effect on the working memory of a visual stimulus with and influence on lick responses more than a second later. dSPNs stimulation changed the cortical activity pattern during the delay to make it more similar to a memory of a stimulus presented contralateral to the stimulated hemisphere (see also Sippy et al., 2015). The activation of iSPNs changed cortical activity in the opposite direction. These results are thereby in support of a recent study demonstrating that the effects of inhibition of the direct and indirect pathways of the dorsomedial striatum are most pronounced in tasks with a memory requirement (Bolkan et al., 2022). Interestingly, the striatum appears to not only influence working memories but also the distribution of attention. Wang et al. (2018) trained mice to direct their attention to one of two stimuli in a task in a change detection task. dSPN activation caused a shift of attention to the stimulus in the contralateral hemifield. Attended stimuli are more likely to be remembered than non-attended ones (Chun and Turk-Browne, 2007; Reeves and Sperling, 1986), and it is therefore probable that overlapping circuits of the basal ganglia contribute to attention shifts and working memory.

Our result extent recent findings that the ventromedial part of the thalamus and the substantia nigra pars reticulata, part of the cortico-striatal loop, are causally involved in working memory. In a task in which mice judged the amplitude of a whisker stimulus, unilateral activation and deactivation of these nuclei during a memory delay had opposite effects on lick-direction (Wang et al., 2021). Here we observed opposite effects of dSPNs and iSPNs in a task in which mice memorized a visual stimulus. Indeed, sustained activity can be found in the delay period of working memory tasks in all areas involved in this loop (this study and e.g. Chen et al., 2017; Goard et al., 2016b; Lee et al., 2020; Wang and Krauzlis, 2020; Wang et al., 2018). One exciting possibility is that the direct pathway maintains the memory, because the loop with its two inhibitory connections can cause neurons in a cortical area to provide positive feedback to themselves. If true, the indirect pathway with its three inhibitory connections could play a role in forgetting working memories that are no longer needed, by decreasing the activity level in the feedback loop of the direct pathway. A study by Tecuapetla et al. (2016) provided some evidence for this view. They trained mice to make a series of lever presses for a food reward. Stimulation of dSPNs caused, under some conditions, an increase in the number of presses. prolonging the motor program. Stimulation of iSPNs, on the other hand, caused the mice to abort the task and start doing something else, as if they completely forgot the motor program in which they were engaged.

The present results support that immediate and delayed visuomotor transformations are, in part, mediated by the loop between the cortex and the basal ganglia, although the loops from cortex to cerebellum to the thalamus and then back to the cortex may also play a role (Gao et al., 2019). Persistent neuronal activity related to working memories appear to be an emergent property of distributed networks (Christophel et al., 2017; Voitov and Mrsic-flogel, 2022) where the striatal pathways can orchestrate cortical activity by integrating sensory and motivational inputs to support adaptive behavior.

## Materials and Methods

### Subjects

We included 6 Thy1-5.17 GCaMP6f mice (Dana et al., 2014), 8 D1-cre, 10 D2-cre, 9 Thy1-5.17 GcaMP6f X D1-cre, and 7 Thy1-5.17 GcaMP6f X Drd2-cre of mixed gender, aged 2-6 months at the start of the experiment (**Table S2**). Animals were housed either in pairs or in isolation and kept on a 12 hour/12 hour reversed day/night cycle. Experiments were performed in the dark phase. All experimental procedures complied with the National Institutes of Health Guide for Care and Use of Laboratory Animals and the study protocol was approved by the ethical committee of the Royal Netherlands Academy of Arts and Sciences and the CCD. The experiments were performed in accordance with all relevant guidelines and regulations.

### Surgery – preparation

Anesthesia was induced using 3-5% isoflurane in oxygen enriched air (50% air, 50% O_2_) in an induction box. The mice were positioned in a stereotactic frame and the depth of anesthesia was monitored throughout the surgery by frequently checking paw reflexes and breathing rate and the concentration of isoflurane was adapted accordingly (between 0.8-2.5%). We subcutaneously injected 5mg/kg meloxicam (0.5 mg/ml) as general analgesic. We monitored the temperature of the animal and kept it between 36.5° and 37.5° with a heating pad coupled to a rectal thermometer. Eyes were covered with ointment to prevent dehydration. The area of incision was shaved, cleaned with alcohol and betadine or hibicet, and lidocaine spray was applied to the skin as local analgesic. An incision was made in the skin along the anteroposterior midline, exposing the skull above the cortex and posterior to lambda. The periost and other tissue was removed from the skull by scraping, rinsing and briefly applying H_2_O_2_ or hibicet.

### Surgery – clear skull and head plate

Once the dorsal skull was completely exposed and dry, a thin layer of adhesive (cyanoacrylate glue Bison) was applied to the bone, thereby making the bone transparent. This effect occurs over the course of the following days and is referred to as the “clear skull cap” technique (Guo et al., 2014b). A thin layer of clear dental cement (C&B super-bond) and nail polish (Electron Microscopy Sciences) were applied for strengthening and to reduce light glare during imaging. A platform of dental cement (Heraeus Charisma) was built posterior to lambda to place the head-bar (for head-fixation purposes). Multiple layers of cement were used to secure the head-bar on the skull. On the outer edges of the clear skull, a small wall of cement (Heraeus Charisma) was built to prevent the skin from growing over the area of interest. The mice were monitored and kept warm while recovering from anesthesia. The mice had a minimum of two days to recover before they were habituated to set-ups and trained.

### Surgery – optogenetics

In D1/2-cre positive mice we additionally performed two craniotomies. The first was for the injection of the virus in the right hemisphere, at 2.5 ML, +0.14 AP from Bregma. A glass pipette with AAV5-Syn-Flex-rcChrimsonR-tdTomato (Addgene 62723, 160nl in total, 1.1*10^13 GC/ml, diluted 1:0.5-2 in saline) or AAV1 EF1a DIO hChR2(H134R) eYFP WPRE hGH (Addgene 20298, 160nl in total, 2.216*10^13 GC/ml, diluted 1:0.5-2 in saline) was lowered to a depth of -4mm relative to Bregma. A total of 160nl was injected in 8 pulses of 10nl/sec, with 30 seconds in between pulses. The second craniotomy was made over the left hemisphere, 2mm lateral and 0.14 anterior to Bregma. A custom-made glass optical fiber (200micron diameter, 0.39NA, ∼5.5mm long) was inserted in an angle of 48 degrees, such that the tip of the glass-fiber would end ∼1mm above the center of virus injection. The ferrule was secured with dental cement (Heraeus Charisma and C&B super-bond). After securing the ferrule the clear-skull procedure was applied.

### Visual stimuli

Visual stimuli were created using the Cogent toolbox (developed by John Romaya at the LON at the Wellcome department of Imaging Neuroscience) and the luminance profile of the monitor was linearized. Some mice included in **Figure 3B** were positioned 11cm (from the eye) in front of a 24-inch LCD monitor (1920 × 1200 pixels, Dell U2412M) (**Table S2**). All other experiments were done in the wide-field imaging set-up with a different LCD monitor (122 × 68cm, liyama LE5564S-B1). These mice were positioned at a distance of 14cm from the screen. We applied a previously described correction for the larger distance between the screen and the mouse at higher eccentricities (Marshel et al., 2011). This method defines stimuli on a sphere and calculates the projection onto a flat surface. Figures were comprised of 100% contrast sinusoidal gratings, with a diameter of 35°, 0.08 cycles/° and mean luminance of 20cd/m^2^. The gratings appeared on the screen at an eccentricity of 40°and 15° elevated relative to the nose of the mouse. Oriented figures (45° for figures on the left side and 135° for figures on the right side) were presented on a grey background (20cd/m^2^), moving with a speed of 24deg/s in a direction orthogonal to the orientation.

### Training mice on the behavioral tasks

We handled the mice daily for 5-10 minutes before we started the training protocol (Guo et al., 2014a). Head-fixation training started by holding the head-bar for a few seconds in the home cage. After one or two days, the animals were head-fixed daily for an increasing amount of time, until they were accustomed to being head-fixed on the set-up. At this point, the animals were put on a fluid restriction protocol with a minimal intake of 0.025ml/g per day, while their health was carefully monitored. First, animals (except Thy1-GcaMP animals, which were only trained on the two-alternative forced choice task, described below) were trained to indicate the appearance of a visual stimulus (visual detection task) by licking either side of a custom-made double lick-spout (**Figure 3A**). Licks were registered by measuring a change in capacitance with an Arduino using custom-written software. A lick to either side of the lick spout was counted as correct and rewarded with 5-8μl of water or milk (Nutrilon). Stimuli on the left were followed by fluid delivery on the left and stimuli on the right by fluid delivery on the right. To prevent mice from licking continuously, they had to withhold licking for a period of 2-4 seconds (uniform distribution) prior to the start of a trial.

Some animals received additional training in a two-alternative-forced-choice (2AFC) task after they completed the visual detection task. In this task they had to indicate the side on which a figure appeared by licking the corresponding side of the double lick-spout. A trial started after a variable inter trial interval (ITI) of 6-10 seconds when the stimulus appeared on the screen for 500ms. Now correct responses required the first lick on the side of the visual stimulus. If the animal made an error, a 5s timeout was added to the ITI. We gradually increased the task-difficulty by increasing the delay between stimulus and response from 0 to 1500ms over several weeks of training, based on performance (staircase). In the final working memory task, mice had to withhold a lick-response for 2000ms (including the 500ms stimulus time). The delay was enforced by attaching the lick-spout to a servo motor (Arduino). The servo motor moved the lick-spout to a position that mice were still able to reach, but at a distance that was not comfortable. Most mice quickly learned that licking the spout at this distance did not yield reward. Other mice persevered premature licking behavior even with the lick-spout at some distance. For these animals we aborted trials immediately after premature lick-responses to discourage this behavior. A time-out of five seconds followed incorrect lick-responses.

### Wide field imaging

After habituation to the set-up, mice were placed under a wide-field fluorescence microscope (Axio Zoom.V16 Zeiss/Caenotec) to image a large part of the cortical surface. Images were captured at 20Hz by a high-speed sCMOS camera (pco.edge 5.5) and recorded using the Encephalos software package (Caenotec). We monitored size and position of the right pupil (50-100Hz sampling rate) and movements of the mouse with a piezo plate under the front paws (100Hz sampling rate).

### Optogenetics

For optogenetic stimulation of the striatum we used a fiber-coupled DPSS Laser (Shanghai Lasers & Optics Century Co.) emitting blue light (BL473T3-100FC, wavelength 473nm) for mice injected with ChR2, and red light (RLM638TA, wavelength 638nm) for mice injected with ChrimsonR. Based on data from pilot experiments, we used 15Hz stimulation with a 10ms pulse width. The duration of the light train differed between experiments. During spontaneous behavior we stimulated for 1 second. In the visual detection task optogenetic stimulation occurred in 40% of trials. In 20% of trials, stimulation occurred 0.5s prior to visual stimulus onset, and in the other 20% of trials it started after the first lick following visual stimulus onset. In the working memory task we stimulated for 500ms. The time between two successive optogenetic stimulations was at least 5 seconds, but usually between 8 seconds to a few minutes. Optogenetic stimulation during the working memory task only occurred when the accuracy of the mice was above 65% in the preceding 15 trials, to prevent the development of lasting behavioral biases as a result of frequent unilateral optogenetic stimulation.

Light intensity was calibrated per mouse: prior to implantation the throughput of the optic fibers was measured with a standard photodiode power sensor (S120C Thorlabs). Light intensity was usually around 2.5mW/mm^2^, but increased gradually to maximally 5mW/mm^2^ if no effect of optogenetic stimulation was observed. These light levels do not cause measurable heating, especially if the light is delivered in pulses of 10ms (Owen et al., 2019; Stujenske et al., 2015). To remove the possibility that the light itself was perceived as a cue, the fiber was within the light-shield of the mouse that was also used for widefield imaging. In addition, a LED positioned close to the fiber but outside the light shield flashed light with a similar wavelength at random times, but with identical duration, pulse width and frequency, to prevent that mice could use optogenetic stimulation as a cue to adapt their behavior.

### Pre-processing of wide-field imaging data

Images were recorded at 20Hz (50ms exposure) and stored in 12-bit, 1600×1600 pixel images (∼15µm per pixel). Images were binned into 800×800 pixels and converted to 16-bit. For each session, images were semi-automatically registered (using multi-modal intensity-based translation, rotation and scaling) to a population receptive field mapping imaging session (van Beest et al., 2021). After registration, images were smoothed using a Gaussian filter with a standard deviation of 2 pixels, and stored in a data-matrix of 400 × 400 × time × number of trials. A top-view of the Allen Brain common coordinate framework was fit to the pRF-map (Allen Institute for Brain Science, 2017). We computed the average ΔF/F (relative to the baseline fluorescence in a 300ms window before stimulus onset). We only included trials in which the accuracy of the mouse was higher than 55% for both left and right stimuli, in a window of 50 trials

When averaging data across mice for top brain views (e.g. beta-weights in **Figure 4F**), each mouse’s fit to the Allen Brain common coordinate framework was aligned to one ‘template’ mouse, using an affine 2D-transformation. For results shown across brain areas (e.g. time-courses, statistics) we first averaged across pixels in each region for individual mice, based on the fit with the Allen Brain atlas before averaging across mice.

### Statistical Analysis

To analyze licking behavior, we not only used lick count (the absolute number of licks in the given time window), but also calculated the lick switch index which is defined as in Equation 1 (above) (**Figure S2B**).

To analyze the imaging data, we calculated neuronal d-prime, which is a measure of how well activity (i.e. ΔF/F) differentiates between two different conditions (e.g. optogenetic stimulation versus baseline activity) on single trials.

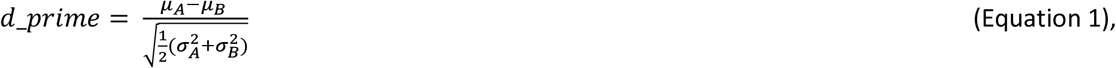

Here *µ*_*A*_ is the average activity for condition A, *µ*_*B*_ the average activity for condition B, and 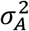 and 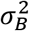 the variance over trials from condition A and B, respectively.

For the memory task we calculated the accuracy, which was defined as:

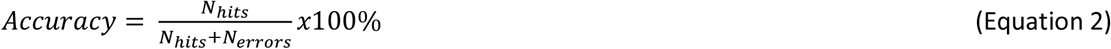

and the omission percentage, which was defined as:

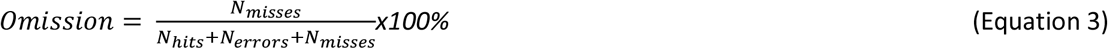

We used build-in MATLAB functions to perform ANOVAs. We used mixed-effect models in the form of

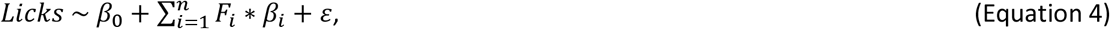

in which *β*_*i*_ is an estimate of how much each factor (*F*_*i*_) contributes to the number of *Licks*. To find the best estimates (minimizing the error *ε*) we used MATLAB’s function fitglme. The identity of the mouse was taken into account as a random factor (i.e. random offsets for individual mice in the mixed-effect model). Mixed-effect models were also applied to neural data (i.e. predicting ΔF/F). We used the publicly available Matlab Toolbox emmeans (J. Hartman, 2019, https://github.com/jackatta/estimated-marginal-means) for post-hoc Wald tests of coefficients. To obtain the significance level of the different factors, model outputs were evaluated with an ANOVA (in Matlab).

### Decoding strategies

We applied multi-pixel pattern decoding strategies to determine how well brain activity could be used to decode the side of the visual stimulus and the lick response. Decoding accuracy was calculated by comparing the predicted label to the actual labels. To prevent biasing the decoding models, we selected an equal number of trials from each condition, 25% of correct and 25% error trials with stimulus_ContraOpto_ and 25% of correct and 25% error trials with stimulus_IpsiOpto_. We used five-fold cross-validation, training the model on 80 percent of data to predict the labels of the other 20 percent. We used the “least dirty” method from the multi-task learning for structural regularization (MALSAR) toolbox for MATLAB (Zhou et al., 2012). MALSAR assigned weights to as few pixels as possible to decode the side of the stimulus and the lick response. We ran the model on data from different epochs in a trial, for individual mice. In some instances, we trained the model on data from a certain condition (e.g. optogenetic stimulation OFF) and tested the model on data from another condition (e.g. early optogenetic stimulation), e.g. in the analysis of **Figure 5D**. To establish significance of the predictions and beta-values for individual mice, we performed bootstrapping (i.e. repeated the procedure 1,000 times while randomizing trial labels with replacement). The significance was determined as the percentile of the prediction in the distribution obtained by bootstrapping.

### Histology

To examine virus expression, we euthanized mice with nembutal and transcardially perfused them with phosphate buffered saline (PBS) followed by 4% paraformaldehyde (PFA) in PBS. We extracted the brain and post-fixated it overnight in 4% PFA before moving it to a PBS solution. We cut the brains into 50µm thick coronal slices and mounted them on glass slides. We imaged the slices on a Zeiss Axioscan Z1 or Zeiss Axioplan 2 microscope (10x objective, Zeiss plan-apochromat, 0.16NA) using custom written Image-Pro Plus software and aligned the images in 3D to the Allen Brain common coordinate framework using a slightly adapted version of a publicly available toolbox (http://github.com/petersaj/AP_histology). We visualized the spread of the virus and determined the optic fiber tract in 3D for individual mice. We modelled the spread of the light from the fiber using software described in Stujenske et al. (2015) and only included mice with virus expression in the striatum and an appropriate placement of the fiber in the D1 and D2 groups. We excluded mice with virus expression that was too widespread. We included mice with no (visible) expression or expression in areas that were not reached by the light as controls. Average intensity maps of expression across D1 and D2 mice were generated by thresholding normalized expression levels (values larger than >99% of all values across all channels) for individual mice, and projecting this binary expression around Bregma +0.14AP±550micron on a 3D aligned coronal slice at Bregma +0.14AP. The intensity level of a given pixel of the coronal slice at Bregma +0.14 (**Figure 1E**) represents the number of mice for which that pixel had expression levels above this threshold. Fiber paths were drawn in 3D for individual mice and projected in a similar way.

## Acknowledgements

We thank the animal and mechatronics departments at the Netherlands Institute for Neuroscience for their assistance. We thank Areg Barsegyan, Ulf Schnabel, Thijs Baaijen and Cédric Gillissen for their assistance in setting up widefield imaging and analysis. We thank Roxana Kooijmans for advice on histology, Emma Ruimschotel and Christiaan Levelt for support with breeding, Chris van der Togt, Mustafa Hamada, Ralph Hamelink and Nicole Yee for technical assistance, Kor Brandsma, Bastijn van den Boom and Sreedeep Mukherjee for surgical assistance and advice and Rudolf Faust, Chris Klink and Lisa Kirchberger for help and advice throughout the experiments. The work was supported by the European Union’s Horizon 2020 Research and Innovation Program (Framework Partnership Agreement No. 650003 Human Brain Project).

## Author Contributions

EvB developed the behavioral tasks and set-up combined widefield imaging and optogenetics. EvB, with help from CM, performed the visual detection experiments combined with optogenetics. EvB, with help from CB, performed the working memory experiments in control mice. EvB, with help from MAOM, LC and BP performed the combined widefield and optogenetic experiments. EvB analyzed the data with input from MWS and PRR. EvB, with help from CM, CB, LC and BP performed histology. EvB, PRR and IW conceived of the experiments, and PR, IW and MWS supervised the project. EvB wrote the paper with help from MWS, IW and PRR.

## Data and materials availability

All data and the computer code used to analyze the data will be made available for download and curated at the Human Brain Project Joint Platform. For clarification, please contact EvB. Correspondence and requests for materials can be sent to PRR.

## References

Alexander, G.E., DeLong, M.R., and Strick, P.L. (1986). Parallel organization of functionally segregated circuits linking basal ganglia and cortex. Annu. Rev. Neurosci. VOL. 9, 357–381.

Allen Institute for Brain Science (2017). Allen Mouse Brain Connectivity Atlas.

Baddeley, A. (1992). Working memory. Science (80-.). 255, 556–559.

van Beest, E.H., Mukherjee, S., Kirchberger, L., Schnabel, U.H., van der Togt, C., Teeuwen, R.R.M., Barsegyan, A., Meyer, A.F., Poort, J., Roelfsema, P.R., et al. (2021). Mouse visual cortex contains a region of enhanced spatial resolution. Nat. Commun. DOI: 10.1038/s41467-021-24311-5.

Bolkan, S., Stone, I., Pinto, L., Ashwood, Z., Garcia, J., Herman, A., Singh, P., Bandi, A., Cox, J., Zimmerman, C., et al. (2022). Opponent control of behavior by dorsomedial striatal pathways depends on task demands and internal state. Nat. Neurosci. 25, 2021.07.23.453573.

Chen, T., Li, N., Daie, K., and Svoboda, K. (2017). A Map of Anticipatory Activity in Mouse Motor Cortex. Neuron 94, 866–879.

Chernysheva, M., Sych, Y., Fomins, A., Luis, J., Warren, A., Lewis, C., Capdevila, L.S., Boehringer, R., Amadei, E.A., Grewe, B.F., et al. (2021). Striatum-projecting prefrontal cortex neurons support working memory maintenance. BioRxiv 1–38.

Choi, E.Y., Thomas Yeo, B.T., and Buckner, R.L. (2012). The organization of the human striatum estimated by intrinsic functional connectivity. J. Neurophysiol. 108, 2242–2263.

Christophel, T.B., Klink, P.C., Spitzer, B., Roelfsema, P.R., and Haynes, J.-D. (2017). The Distributed Nature of Working Memory. Trends Cogn. Sci. 21, 111–124.

Chun, M.M., and Turk-Browne, N.B. (2007). Interactions between attention and memory. Curr. Opin. Neurobiol. 17, 177–184.

Cox, J., and Witten, I.B. (2019). Striatal circuits for reward learning and decision-making. Nat. Rev. Neurosci. DOI: 10.1038/s41583-019-0189-2.

Cruz, B.F., Guiomar, G., Soares, S., Motiwala, A., Machens, C.K., and Paton, J.J. (2022). Action suppression reveals opponent parallel control via striatal circuits. DOI: 10.1038/s41586-022-04894-9.

Curtis, C.E., and D’Esposito, M. (2003). Persistent activity in the prefrontal cortex during working memory. Trends Cogn. Sci. 7, 415–423.

Dana, H., Chen, T.-W., Hu, A., Shields, B.C., Guo, C., Looger, L.L., Kim, D.S., and Svoboda, K. (2014). Thy1-GCaMP6 Transgenic Mice for Neuronal Population Imaging In Vivo. PLoS One 9, e108697.

Ding, L., and Gold, J.I. (2010). Caudate encodes multiple computations for perceptual decisions. J. Neurosci. 30, 15747–15759.

Dotson, N.M., Hoffman, S.J., Goodell, B., and Gray, C.M. (2018). Feature-Based Visual Short-Term Memory Is Widely Distributed and Hierarchically Organized. Neuron 1–12.

Esmaeili, V., Tamura, K., Muscinelli, S.P., Modirshanechi, A., Boscaglia, M., Lee, A.B., Oryshchuk, A., Foustoukos, G., Liu, Y., Crochet, S., et al. (2021). Rapid suppression and sustained activation of distinct cortical regions for a delayed sensory-triggered motor response. Neuron 109, 2183-2201.e9.

Foster, N.N., Barry, J., Korobkova, L., Garcia, L., Gao, L., Becerra, M., Sherafat, Y., Peng, B., Li, X., Choi, J.-H., et al. (2021). The mouse cortico–basal ganglia–thalamic network. Nature 598, 188–194.

Frank, M.J., Loughry, B., and O’Reilly, R.C. (2001). Interactions between frontal cortex and basal ganglia in working memory: a computational model. Cogn. Affect. Behav. Neurosci. 1, 137–160.

Fuster, J.M., and Alexander, G.E. (1971). Neuron activity related to short-term memory. Science (80-.). 1–4.

Fuster, J.M., and Alexander, G.E. (1973). Firing changes in cells of the nucleus medialis dorsalis associated with delayed response behavior. Brain Res. 61, 79–91.

Fuster, J.M., and Jervey, J.P. (1981). Inferotemporal neurons distinguish and retain behaviorally relevant features of visual stimuli. Science (80-.). 212, 952–955.

Gao, Z., Davis, C., Thomas, A.M., Economo, M.N., Abrego, A.M., Svoboda, K., De Zeeuw, C.I., and Li, N. (2019). A cortico-cerebellar loop for motor planning. Nature 563, 113–116.

Goard, M.J., Pho, G.N., Woodson, J., and Sur, M. (2016). Distinct roles of visual, parietal, and frontal motor cortices in memory-guided sensorimotor decisions. Elife 5, 1–30.

Graybiel, A.M. (1998). The Basal Ganglia and Chunking of Action Repertoirs. Neurobiol. Learn. Mem. 70, 119–136.

Groenewegen, H.J. (2003). The basal ganglia and motor control. Neural Plast. 10, 107–120.

Guo, Z. V., Hires, S.A., Li, N., O’Connor, D.H., Komiyama, T., Ophir, E., Huber, D., Bonardi, C., Morandell, K., Gutnisky, D., et al. (2014a). Procedures for Behavioral Experiments in Head-Fixed Mice. PLoS One 9, e88678.

Guo, Z. V., Inagaki, H.K., Daie, K., Druckmann, S., Gerfen, C.R., and Svoboda, K. (2017). Maintenance of persistent activity in a frontal thalamocortical loop. Nature 545, 181–186.

Guo, Z. V, Li, N., Huber, D., Ophir, E., Gutnisky, D., Ting, J.T., Feng, G., and Svoboda, K. (2014b). Flow of cortical activity underlying a tactile decision in mice. Neuron 81, 179–194.

Haber, S.N. (2016). Corticostriatal circuitry. Dialogues Clin. Neurosci. 18, 7–21.

Hikosaka, O., and Wurtz, R.H. (1983). Visual and oculomotor functions of monkey substantia nigra pars reticulata. I. Relation of visual and auditory responses to saccades. J. Neurophysiol. 49, 1230–1253.

Hikosaka, O., Sakamoto, M., and Usui, S. (1989). Functional properties of monkey caudate neurons. III. Activities related to expectation of target and reward. J. Neurophysiol. 61, 814–832.

Van Kerkoerle, T., Self, M.W., and Roelfsema, P.R. (2017). Layer-specificity in the effects of attention and working memory on activity in primary visual cortex. Nat. Commun. 8, 1–12.

Kubota, K., and Niki, H. (1971). Prefrontal cortical unit activity and delayed alternation performance in monkeys. J. Neurophysiol. 34, 337–347.

Lee, J., and Sabatini, B.L. (2021). Striatal indirect pathway mediates exploration via collicular competition. Nature DOI: 10.1038/s41586-021-04055-4.

Lee, J., Wang, W., and Sabatini, B.L. (2020). Anatomically segregated basal ganglia pathways allow parallel behavioral modulation. Nat. Neurosci. DOI: 10.1038/s41593-020-00712-5.

Lee, J.Y., Jun, H., Soma, S., Nakazono, T., Shiraiwa, K., Dasgupta, A., Nakagawa, T., Xie, J.L., Chavez, J., Romo, R., et al. (2021). Dopamine facilitates associative memory encoding in the entorhinal cortex. Nature DOI: 10.1038/s41586-021-03948-8.

Marshel, J.H., Garrett, M.E., Nauhaus, I., and Callaway, E.M. (2011). Functional specialization of seven mouse visual cortical areas. Neuron 72, 1040–1054.

McGeorge, a J., and Faull, R.L. (1989). The organization of the projection from the cerebral cortex to the striatum in the rat. Neuroscience 29, 503–537.

Mendoza-Halliday, D., Torres, S., and Martinez-Trujillo, J.C. (2014). Sharp emergence of feature-selective sustained activity along the dorsal visual pathway. Nat. Neurosci. 17, 1255–1262.

Mushiake, H., and Strick, P.L. (1995). Pallidal neuron activity during sequential arm movements. J. Neurophysiol. 74, 2754–2758.

Nonomura, S., Nishizawa, K., Sakai, Y., Kawaguchi, Y., Kato, S., Uchigashima, M., Watanabe, M., Yamanaka, K., Enomoto, K., Chiken, S., et al. (2018). Monitoring and Updating of Action Selection for Goal-Directed Behavior through the Striatal Direct and Indirect Pathways. Neuron 99, 1302-1314.e5.

Oldenburg, I.A.A., and Sabatini, B.L.L. (2015). Antagonistic but Not Symmetric Regulation of Primary Motor Cortex by Basal Ganglia Direct and Indirect Pathways. Neuron 86, 1174–1181.

Orsolic, I., Rio, M., Mrsic-Flogel, T.D., and Znamenskiy, P. (2021). Mesoscale cortical dynamics reflect the interaction of sensory evidence and temporal expectation during perceptual decision-making. Neuron 1–15.

Osaka, N., Logie, R.H., and D’Esposito, M. (2012). The Cognitive Neuroscience of Working Memory. Cogn. Neurosci. Work. Mem. 1–408.

Owen, S.F., Liu, M.H., and Kreitzer, A.C. (2019). Thermal constraints on in vivo optogenetic manipulations. Nat. Neurosci. 22, 1061–1065.

Peixoto, D., Verhein, J.R., Kiani, R., Kao, J.C., Nuyujukian, P., Chandrasekaran, C., Brown, J., Fong, S., Ryu, S.I., Shenoy, K. V., et al. (2021). Decoding and perturbing decision states in real time. Nature 591, 604–609.

Peters, A.J., Fabre, J.M.J., Steinmetz, N.A., Harris, K.D., and Carandini, M. (2021). Striatal activity reflects cortical activity patterns. Nature DOI: 10.1038/s41586-020-03166-8.

Rainer, G., Rao, S.C., and Miller, E.K. (1999). Prospective coding for objects in primate prefrontal cortex. J. Neurosci. 19, 5493–5505.

Reeves, A., and Sperling, G. (1986). Attention Gating in Short-Term Visual Memory. Psychol. Rev. 93, 180–206.

Ren, C., and Komiyama, T. (2021). Characterizing cortex-wide dynamics with wide-field calcium imaging. J. Neurosci. 41, 4160–4168.

Rosenthal, R., Rubin, D., and Meng, X.-L. (1992). Comparing correlated correlation coefficients. Psychol. Bull. 111, 172–175.

Rossi, M.A., Li, H.E., Lu, D., Kim, I.., Bartholomew, R.A., Gaidis, E., Barter, J., Kim, N., Tong Cai, M., Soderling, S., et al. (2016). A GABAergic nigrotectal pathway for coordination of drinking behavior. Nat. Neuros 19, 742–748.

Saint-Cyr, J.A., Ungerleider, L.G., and Desimone, R. (1990). Organization of visual cortical inputs to the striatum and subsequent outputs to the pallido-nigral complex in the monkey. J. Comp. Neurol. 298, 129–156.

Saunders, A., Oldenburg, I. a, Berezovskii, V.K., Johnson, C. a, Kingery, N.D., Elliott, H.L., Xie, T., Gerfen, C.R., and Sabatini, B.L. (2015). A direct GABAergic output from the basal ganglia to frontal cortex. Nature 05, 85–89.

Schultz, W. (2016). Dopamine reward prediction-error signalling: a two-component response. Nat. Publ. Gr. 17, 183–195.

Schultz, W., Dayan, P., and Montague, P.R. (1997). A neural substrate of prediction and reward. Science (80-.). 275, 1593–1599.

Sippy, T., Lapray, D., Crochet, S., and Petersen, C.C.H. (2015). Cell-Type-Specific Sensorimotor Processing in Striatal Projection Neurons during Goal-Directed Behavior. Neuron 1–8.

Smith, Y., Bevan, M.D., Shink, E., and Bolam, J.P. (1998). Microcircuitry of the direct and indirect pathways of the basal ganglia. Neuroscience 86, 353–387.

Stujenske, J.M., Spellman, T., and Gordon, J.A. (2015). Modeling the Spatiotemporal Dynamics of Light and Heat Propagation for InVivo Optogenetics. Cell Rep. 12, 525–534.

Tecuapetla, F., Jin, X., Lima, S.Q., and Costa, R.M. (2016). Complementary Contributions of Striatal Projection Pathways to Action Initiation and Execution. Cell 166, 703–715.

Voitov, I., and Mrsic-flogel, T.D. (2022). Cortical feedback loops bind distributed representations of working memory. DOI: 10.1038/s41586-022-05014-3.

van Vugt, B., van Kerkoerle, T., Vartak, D., and Roelfsema, P.R. (2020). The contribution of AMPA and NMDA receptors to persistent firing in the dorsolateral prefrontal cortex in working memory. J. Neurosci. 40, 2458–2470.

Wang, X.J. (2001). Synaptic reverberation underlying mnemonic persistent activity. Trends Neurosci. 24, 455–463.

Wang, L., and Krauzlis, R.J. (2020). Involvement of Striatal Direct Pathway in Visual Spatial Attention in Mice. Curr. Biol. 30, 4739-4744.e5.

Wang, L., Rangarajan, K. V, Gerfen, C.R., and Krauzlis, R.J. (2018). Activation of Striatal Neurons Causes a Perceptual Decision Bias during Visual Change Detection in Mice. Neuron 1–13.

Wang, Y., Yin, X., Zhang, Z., Li, J., Zhao, W., and Guo, Z. V. (2021). A cortico-basal ganglia-thalamo-cortical channel underlying short-term memory. Neuron 1–14.

White, N.M. (1997). Mnemonic functions of the basal ganglia. Curr. Opin. Neurobiol. 7, 164–169.

Wise, S.P., Murray, E.A., and Gerfen, C.R. (1996). The frontal cortex-basal ganglia system in primates. Crit. Rev. Neurobiol. DOI: 10.1615/critrevneurobiol.v10.i3-4.30.

Xiong, Q., Znamenskiy, P., and Zador, A.M. (2015). Selective corticostriatal plasticity during acquisition of an auditory discrimination task. Nature DOI: 10.1038/nature14225.

Zhou, J., Chen, J., and Ye, J. (2012). MALSAR: Multi-task Learning via Structural Regularization. Arizona State Univ DOI:.

Znamenskiy, P., and Zador, A.M. (2013). Corticostriatal neurons in auditory cortex drive decisions during auditory discrimination. Nature 497, 482–485.

