## Supplementary Information for "The direct and indirect pathways of the basal ganglia antagonistically influence cortical activity and perceptual decisions"

*Supplementary analysis*

Influence of optogenetic stimulation on the ability of mice to switch lick spout: To investigate whether optogenetic stimulation enforced a specific motor response, we determined the switch index (SI). The SI is defined as:

$$SI = \frac{\sum C \to I - \sum I \to C}{\sum C \to I + \sum I \to C} \quad \text{(Equation 1)}$$

in which $\sum C \to I$ is the number of times the mouse switched from contraversive to ipsiversive licks within a trial, and $\sum I \to C$ the number of times the mouse switched from ipsiversive to contraversive licks within a trial. The SI was positive (i.e. more ipsiversive switches) when the visual stimulus and reward were ipsilateral, and negative (i.e. more contraversive switches) when the visual stimulus and reward were contralateral (repeated measures ANOVA with factors visual stimulus side, genotype and optogenetic stimulation: main effect of visual stimulus side $F_{1,65}=231$, $p<0.001$, **Figure S2B**). Optogenetic stimulation did not change the SI. Hence, the mice switched to the other side in spite of dSPN and iSPN activation indicating that it did not enforce specific lateralized lick responses (no main or interaction effects in the ANOVA; ps>0.05).

Influence of optogenetic stimulation after the first lick in the visual detection task: In the main text we focused our analysis on the condition where optogenetic stimulation started 0.5s prior to the visual stimulus onset. On another 20% of trials, the first lick triggered optogenetic stimulation, which lasted until 1.5s after stimulus onset. We obtained similar, albeit weaker effects compared to the trials with earlier optogenetic stimulation (maximally 0.5 licks difference between no stimulation and stimulation conditions). Given that lick onset times were also variable, we focused our main analysis on trials in which optogenetic stimulation started 0.5s prior to the onset of the visual stimulus.

Correlation analysis of wide-field imaging before and after training in the detection task: To investigate the brain regions most influenced by training, we measured the correlation of the activity of pixels (averaged across all trials) between three conditions, (1) dSPN stimulation in naïve and (2) trained mice, and (3) during the visual detection task (in a time window from 0-0.5s). The correlations were positive ($p<0.001$, **Figure S3C**), showing that if a pixel is active in the task, it is likely to be activated by the optogenetic stimulation as well. Training increased the similarity between the optogenetic effect and the task-related brain activity pattern (comparison of correlations before and after training using Meng's z-test for correlated correlations, Rosenthal et al., 1992, $p<0.001$, **Figure S3C**). Hence, training causes dSPN activation of cortical regions that activate during the task on which the mice are trained.

*Supplementary Figures*

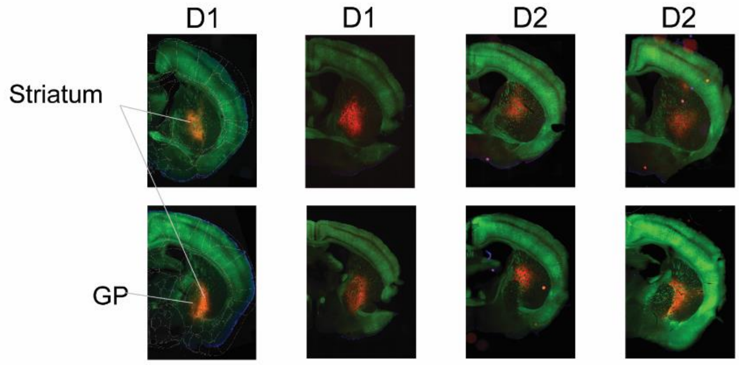


**Figure S1.** Expression of Thy1-GCaMP-GFP (green) and ChrimsonR-tdTomato (red) in example D1/2-cre X Thy1-GCaMP6f mice. Top row is approximately +0.14mm anterior to Bregma, and bottom row 0.4mm posterior to Bregma. ChrimsonR-tdTomato positive cell bodies were mainly found in the striatum.

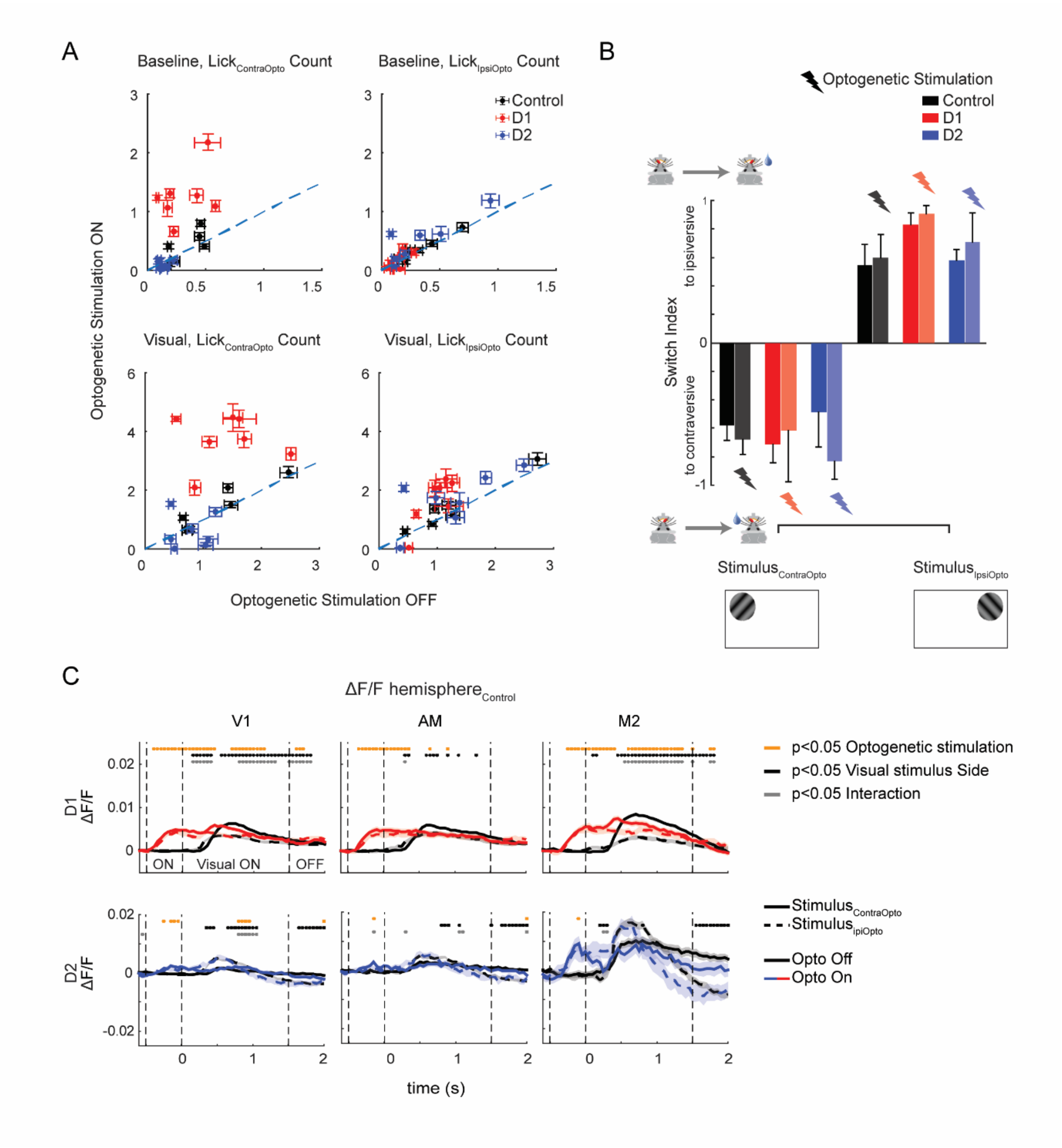


**Figure S2 Effect of optogenetic stimulation of the striatum in a visual detection task.** A) Average lick-count on trials with (x-axis) and without optogenetic stimulation (y-axis). Datapoints represent D1 (red), D2 (blue) and control mice (black). Upper panels, effects in the baseline time window (-500 to 0ms relative to stimulus onset). Lower panels, visual time window (0-1s after stimulus onset). Error bars denote s.e.m. B) Average switch index (SI), which is negative if mice tend to switch to ipsiversive licks (i.e. when reward is delivered on the lick spout at the same side as the optogenetic stimulation) and positive otherwise. As expected, the location of the visual stimulus influenced the switch index (repeated measures ANOVA, $F_{1,65}$=231, p<0.001), but optogenetic stimulation did not. C) Time-courses of GCaMP signal in three example areas in $\text{hemisphere}_{\text{Control}}$ in D1 (N=4) and D2 mice (N=3 mice). Solid (dashed) traces, activity induced by $\text{stimulus}_{\text{ContraOpto}}$ ($\text{stimulus}_{\text{IpsiOpto}}$). The different epochs (pre-stimulus epoch with optogenetic stimulation, visual stimulus on, stimulus off) are indicated by vertical dashed lines. Shaded area denotes s.e.m. The orange dots above indicate a significant main effect of optogenetic stimulation (p<0.05), black dots a main effect of the side of the visual stimulus (p<0.05) and grey dots an interaction between these factors (p<0.05). Note that part of $\text{hemisphere}_{\text{Control}}$ was occluded by the fiber implant.

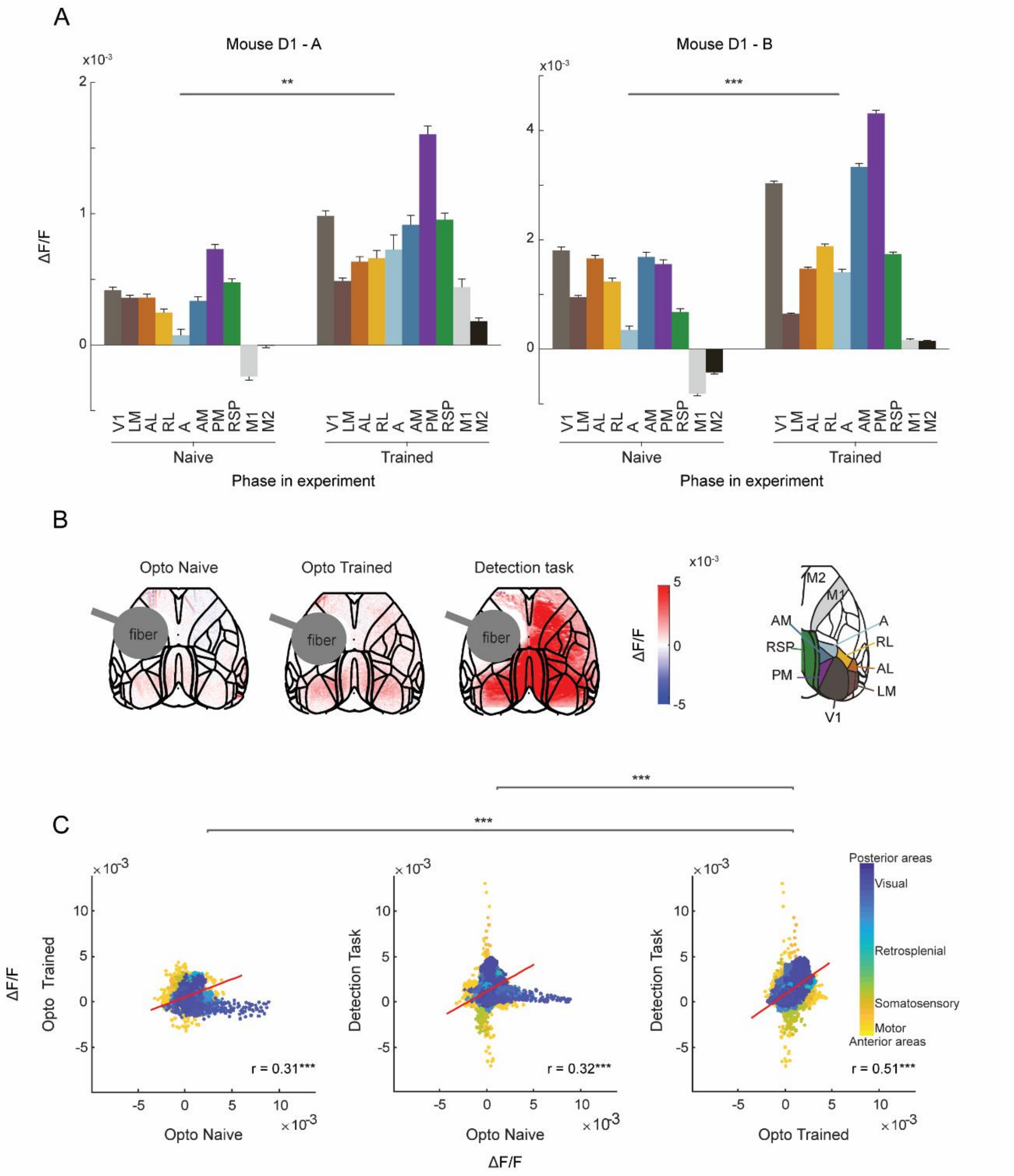


**Figure S3. Effect of dSPN activation in naïve and trained mice outside the task.** A) Activity (ΔF/F) in cortical areas when the striatum was optogenetically stimulated in two D1 mice (mouse A and B) outside the task. Cortical activation evoked by dSPN stimulation increased once the mice had become proficient in the visual detection task (two-way ANOVA with factors area and training revealed a main effect of training: **, p<0.01; ***, p<0.001). B) Influence of dSPN stimulation on cortical activity before (left) and after training (middle) for mouse A. The right panel shows cortical activity for the same mouse during the visual detection task (0-1500ms). C) Correlation of the activity elicited between pairs of the following three conditions: optogenetics outside the task in (1) untrained and (2) trained mice and (3) activity elicited by the task itself (without optogenetic stimulation), across all imaged pixels. Colours denote pixels from different cortical regions (color bar). All pairwise correlations are significant (***,p<001), indicating that if a pixel is active in one of the three conditions, it is likely to be active in the other two. The correlation between activity of pixels during the task and during optogenetics was significantly higher after training than before (Meng's z-test: Z=51.95, p<0.001). Red lines are linear fits. r, correlation coefficient.

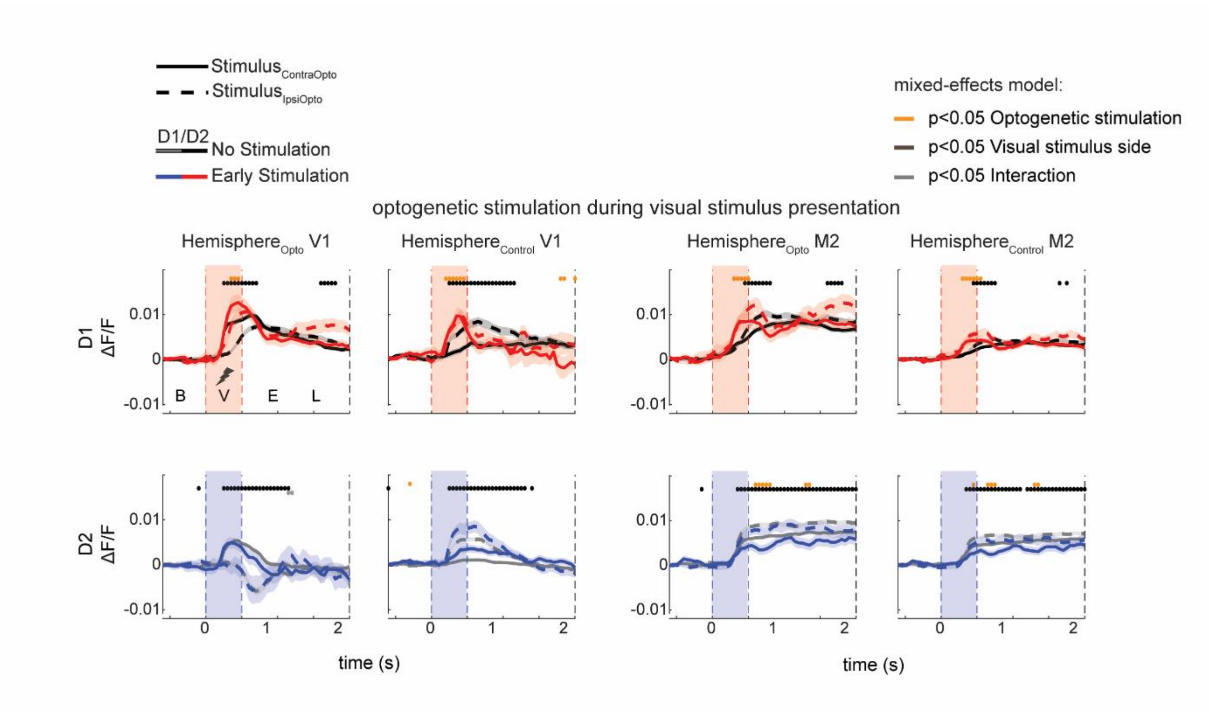


**Figure S4. Optogenetic stimulation of the striatum in the working memory task.** Influence of optogenetic stimulation of the striatum (colored traces) during the presentation of the visual stimulus (0-0.5s after stimulus onset) on activity in M2 and V1. Traces show average $\Delta F/F$ and shaded regions s.e.m. Solid (dashed) lines represent trials with $\text{stimulus}_{\text{ContraOpto}}$ ($\text{stimulus}_{\text{IpsiOpto}}$). Mixed-effects models per time point revealed significant main effects of optogenetic stimulation (orange circles, $p<0.05$), visual stimulus side (black circles, $p<0.05$) and the interaction between these factors (grey circles, $p<0.05$).

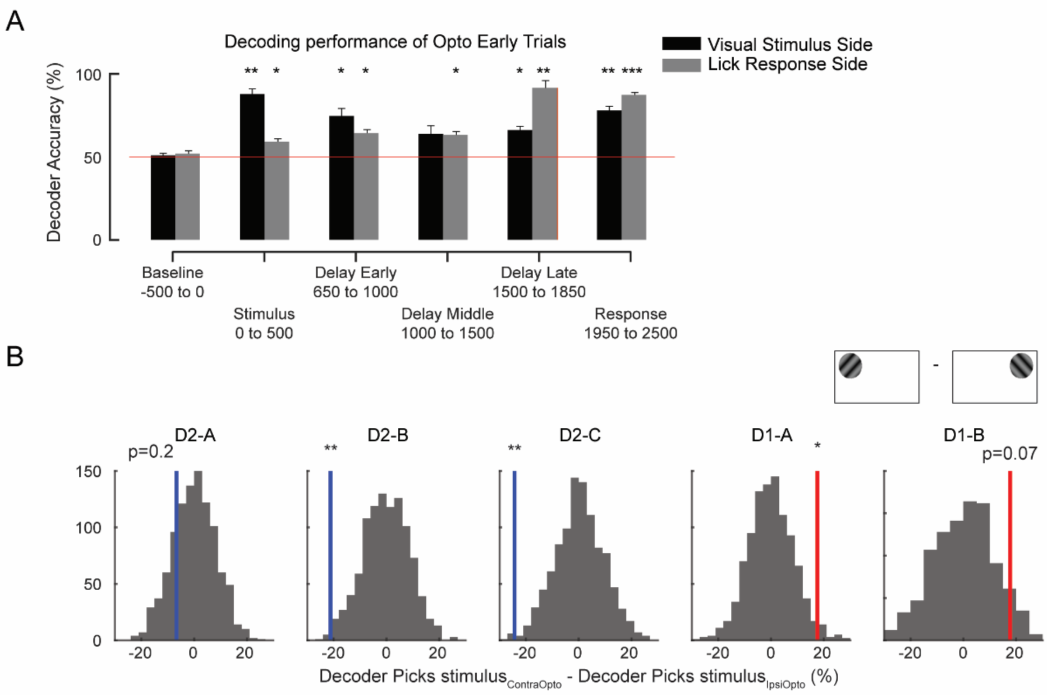


**Figure S5. MALSAR decoding of visual stimulus and lick direction and influence of optogenetic activation.** A) Decoding accuracy of stimulus location side (black bars) and lick direction (grey bars) in D-creXThy1-GCaMp6f mice in trials with early optogenetic stimulation using a model that was trained on trials without optogenetic stimulation. Stars indicate significance; *, p<0.05; **, p<0.01; ***, p<0.001. B) Blue and red lines show the influence of optogenetic stimulation on stimulus decoding in individual D1 and D2 mice. Positive (negative) values indicate an increase in the decoding of $\text{stimulus}_{\text{ContraOpto}}$ ($\text{stimulus}_{\text{IpsiOpto}}$). Histograms show the bootstrapped distributions. *, p<0.05; **, p<0.01.

| Acronym | Full name | Part of |
|---|---|---|
| **V1** | Primary visual cortex | Dorsal cortex |
| **LM** | Lateral visual area | Dorsal cortex |
| **AL** | Anterolateral visual area | Dorsal cortex |
| **RL** | Rostrolateral visual area | Dorsal cortex |
| **A** | Anterior area | Dorsal cortex |
| **AM** | Anteriomedial visual area | Dorsal Cortex |
| **PM** | Posteromedial visual area | Dorsal cortex |
| **RSP** | Retrosplenial area | Dorsal cortex |
| **M1** | Primary motor cortex | Dorsal cortex |
| **M2** | Secondary motor cortex | Dorsal cortex |
| **D1** | Dopamine-1 (Receptor) | Direct pathway basal ganglia |
| **D2** | Dopamine-2 (Receptor) | Indirect pathway basal ganglia |
| **dSPN** | Direct-pathway striatal projection neurons | Basal ganglia (direct pathway) |
| **iSPN** | Indirect-pathway striatal projection neurons | Basal ganglia (indirect pathway) |

Table S1 Acronyms. Brain regions and receptors, their full name and whether they belong to cortex or basal ganglia.

| Mouse | Genotype & Virus | Sex (M/F) | Fig. 1E | Fig. 2 | Fig. 3B | Fig. 3C,D | Fig. 4B,D | Fig. 4C,E,F | Fig. 5 |
|---|---|---|---|---|---|---|---|---|---|
| **1** | D2XChrimsonXGCaMP | F | X | X | | X | | X | X |
| **2** | D1XChrimsonXGCaMP | F | X | X | X | X | | X | X |
| **3** | D2XChrimsonXGCaMP | F | X | X | X | X | | X | X |
| **4** | D1XChrimsonXGCaMP | M | X | X | X | X | | | |
| **5** | D2XChrimsonXGCaMP | M | X | X | X | X | | X | X |
| **6** | D1XChrimsonXGCaMP | F | | X | X | X | | | |
| **7** | D1XChrimsonXGCaMP | F | X | X | X | X | | X | X |
| **8** | D1XChrimsonXGCaMP | F | X | X | | | | | |
| **9** | D2XChr2 | M | X | | X | | | | |
| **10** | D2XChrimson | F | X | | X | | | | |
| **11** | D1XChr2 | M | X | | X | | | | |
| **12** | D1XChr2 | M | X | | X | | | | |
| **13** | D2XChrimsonXGCaMP | M | X | | | | | | |
| **14** | D2XChr2 (control) | M | | | X | | | | |
| **15** | D1XChr2 (control) | M | | | X | | | | |
| **16** | D1XChr2 (control) | M | | | X | | | | |
| **17** | D1XChr2 | M | X | | X | | | | |
| **18** | D2XChr2 | F | X | | X | | | | |
| **19** | D2XChrimson (control) | F | | | X | | | | |
| **20** | D1XChrimson (control) | F | | | X | | | | |
| **21** | D2XChr2 (control) | M | | | X | | | | |
| **22** | D2XChr2 | M | X | | X | | | | |
| **23** | D2XChr2 | M | X | | X | | | | |
| **24** | GCaMP | M | | | | | X | X | |
| **25** | GCaMP | M | | | | | | X | |
| **26** | GCaMP | F | | | | | | X | |
| **27** | GCaMP | F | | | | | | X | |
| **28** | GCaMP | M | | | | | | X | |

Table S2. Mice that contributed to the main results. D1/D2 refers to D1 or D2 – cre positive mice, Chrimson and Chr2 are both DIO-versions, such that the excitatory opsin is only expressed in D1 or D2 cells. GCaMP stands for Thy1-GCaMP6f positive mice. A cell in the table is marked with an X if a mouse contributed to the data in a particular figure. Mice that were excluded from analysis are not shown.
